# Metabolic impact of heterologous protein production in *Pseudomonas putida*: Insights into carbon and energy flux control

**DOI:** 10.1101/2023.07.27.550675

**Authors:** Philippe Vogeleer, Pierre Millard, Ana-Sofia Ortega Arbulú, Katharina Pflüger-Grau, Andreas Kremling, Fabien Létisse

## Abstract

For engineered microorganisms, the production of heterologous proteins that are often useless to host cells represents a burden on resources, which have to be shared with normal cellular processes. Within a certain metabolic leeway, this competitive process has no impact on growth. However, once this leeway, or free capacity, is fully utilized, the extra load becomes a metabolic burden that inhibits cellular processes and triggers a broad cellular response, reducing cell growth and often hindering the production of heterologous proteins. In this study, we sought to characterize the metabolic rearrangements occurring in the central metabolism of *Pseudomonas putida* at different levels of metabolic load. To this end, we constructed a *P. putida* KT2440 strain that expressed two genes encoding fluorescent proteins, one in the genome under constitutive expression to monitor the free capacity, and the other on an inducible plasmid to probe heterologous protein production. We found that metabolic fluxes are considerably reshuffled, especially at the level of periplasmic pathways, as soon as the metabolic load exceeds the free capacity. Heterologous protein production leads to the decoupling of anabolism and catabolism, resulting in large excess energy production relative to the requirements of protein biosynthesis. Finally, heterologous protein production was found to exert a stronger control on carbon fluxes than on energy fluxes, indicating that the flexible nature of *P. putida*’s central metabolic network is solicited to sustain energy production.

**Highlights:** Heterologous protein production in *P. putida* reshuffles the periplasmic metabolism.

Increased protein production progressively decouples catabolism from anabolism.

Protein production exerts a stronger control on energy than on carbon fluxes.

Glucose is directed towards ATP production to meet the elevated energy demands.

## 1. Introduction

*Pseudomonas putida* is widely regarded as a valuable workhorse for white biotechnology due to its combination of rapid growth, minimal nutrient requirements, and versatile metabolism (Nikel and de Lorenzo, 2018). Its resistance to toxic compounds, organic solvents and to variations in pH and temperature also make it attractive for industrial applications. Moreover, *P. putida* lacks the virulence factors commonly found in the genome of other *Pseudomonas* species (*i*.*e*. type III secretion system and endotoxin A) (Udaondo et al., 2016). Its ability to produce endogenous biopolymers (polyhydroxyalkanoates - PHA), biosurfactants (rhamnolipids) and bioplastic synthon (2,5-furandicarboxylic acid - FDCA) is already industrially exploited (Weimer et al., 2020). It is also used in bioremediation processes, as it produces various enzymes that degrade xenobiotic aromatic compounds (de Lorenzo, 2008; Poblete-Castro et al., 2017). This ability to degrade aromatic compounds and its high intrinsic resistance to metabolic and physiological stresses differentiates *P. putida* from other bacterial species used in synthetic biology such as *Escherichia coli* and *Bacillus subtilis* (Nikel et al., 2014), and makes *P. putida* a promising organism for biotechnological applications.

However, biotechnological production is often limited by cellular capacity (Ceroni et al., 2015) and in engineered cells, the heterologous production of proteins represents an unnatural load, which consumes energy and cellular resources, such as ribosomes, polymerases and metabolites. Cells have a certain metabolic leeway, which we call their *free capacity*, that allows them to produce heterologous proteins without reducing their growth rate. Any additional load, however, becomes a burden and the growth rate decreases (Borkowski et al., 2016). The central metabolism responds by providing precursors, and chemical and redox energy for the implemented task, but this also triggers stress-induced mechanisms related to protein production (Wittmann et al., 2007).

*P. putida*’s metabolism is characterized by its particular central carbon network (Figure 1). At the periplasmic level, glucose is primarily directly oxidized to gluconate with a smaller fraction being converted to 2-ketogluconate (2-KG). Once internalized into the cytoplasm by their respective transporters, these metabolites are converted into 6-phosphogluconate (6-PG) either by gluconate kinase (GntK) or *via* the 2-KG bypass, by the activity of 2-KG kinase (KguK) and 2-KG-6-P reductase (KguD) (del Castillo et al., 2007; Vicente and Cánovas, 1973). Furthermore, the classical Embden–Meyerhof–Parnas (EMP) pathway is nonfunctional because of the absence of phosphofructokinase, which normally converts fructose-6-phosphate to fructose-1,6-biP. As a consequence, carbon is processed mainly *via* the Entner-Doudoroff (ED) pathway and partially *via* the pentose phosphate (PP) pathway (Nikel et al., 2015). Compared to the EMP pathway, commonly known as glycolysis, the oxidation of one mole of glucose into two pyruvates through the ED pathway yields less ATP (one mole only) but an equal amount of catabolic and anabolic reducing power (one NADH and one NADPH molecule, respectively) instead of two NADHs. Remarkably, NADPH formation is also boosted by the recycling of the glyceraldehyde-3-phosphate (GAP) formed from glucose in the ED pathway to hexose-phosphate, and the combined action of enzymes from the ED pathway, the EMP pathway (in the gluconeogenic direction) and the PP pathway, in what is collectively known as the EDEMP cycle (Kohlstedt and Wittmann, 2019; Nikel et al., 2015). The EDEMP cycle enables *P. putida* to adjust NADPH production to meet anabolic demands and provides the intrinsic resistance to oxidative stress (Kim and Park, 2014). The cyclic architecture of *P. putida*’s metabolism thus supports stress resistance alongside oxidative energy production. In the context of biotechnology applications (Volke et al., 2020), an important question is how the specific features of *P. putida*’s central metabolism allow the bacteria to cope with the metabolic load incurred by protein overproduction.

**Fig 1.**
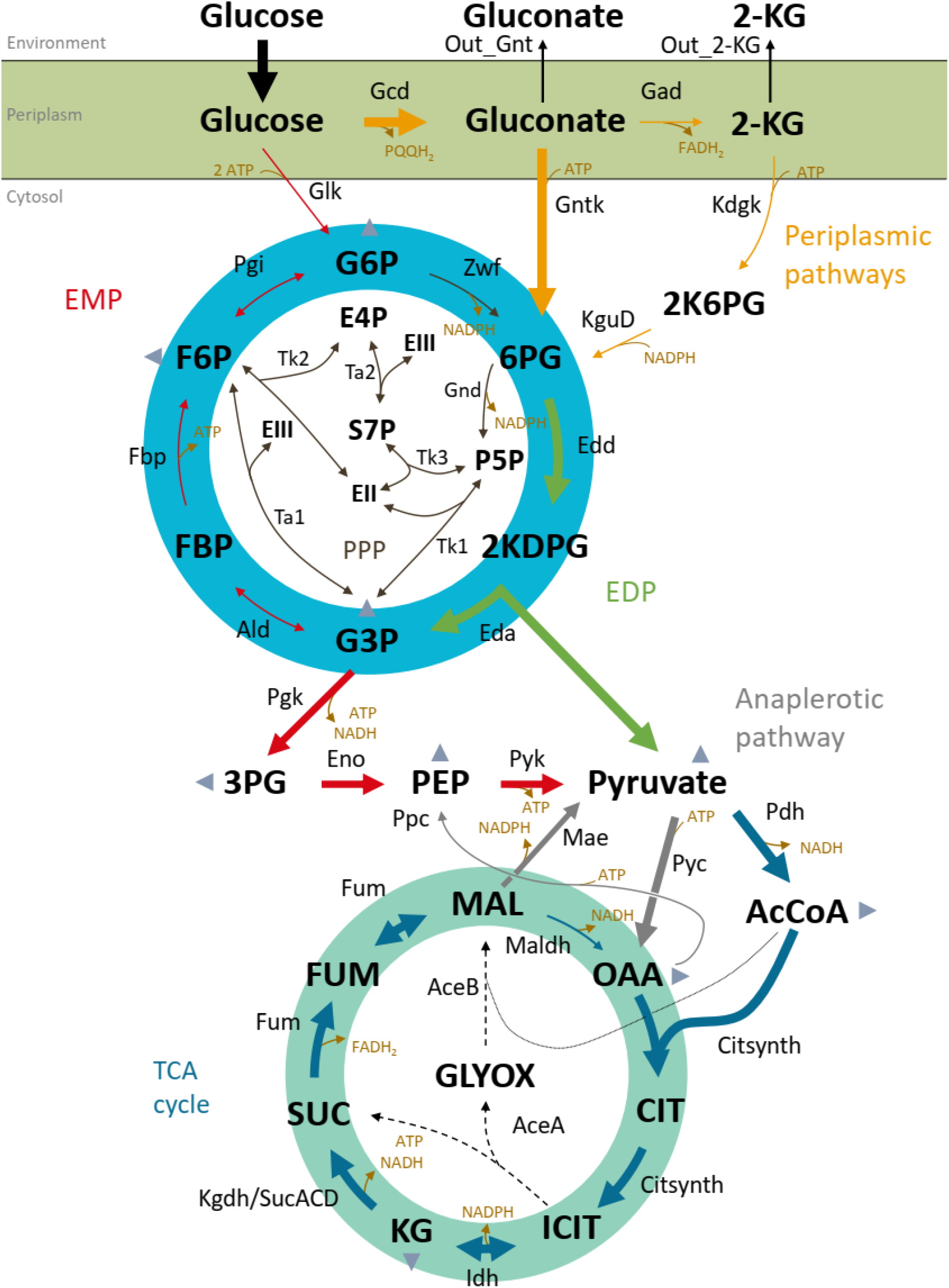
Metabolic pathway of *P. putida* KT2440’s central metabolism. The EDEMP pathway is circled in turquoise and the TCA cycle in water-green. Biomass precursors are indicated by grey triangles. Arrow sizes reflect the intracellular carbon flux of glucose-grown cells relative to the glucose uptake rate (set to 100 %) determined by Kohlstedt and Wittmann (Kohlstedt and Wittmann, 2019). EII and EIII represent the C_2_ and C_3_ fragment pools, respectively, bound to transketolase and transaldolase, and transferred to an aldose acceptor as described by Kleijn et al. (2005). Reaction names and equations are provided in the supplemental data (Table S1).

The aim of this study was to characterize the rearrangements in *P. putida*’s central metabolism at different levels of protein overproduction. Following Ceroni *et al*. (2015), we constructed a *P. putida* KT2440 strain that i) constitutively expressed the *mCherry* gene to monitor cellular capacity and ii) carried a plasmid encoding maltose-binding protein (MBP) fused to fluorescent eGFP for better detection under the control of an inducible promoter. Protein production was tuned by varying the concentrations of the inducer, and quantified by measuring the level of green fluorescence. The metabolic response to the different levels of metabolic burden induced by gradually increasing heterologous protein production was investigated by measuring bacterial growth and carbon and energy fluxes. Comparisons between metabolic flux maps obtained under induced and non-induced conditions provide insights into the flux rearrangements within the central metabolism caused by protein production.

## 2. Material and Methods

### 2.1 Strains and plasmids

*P. putida* and *E. coli* strains were cultivated at 30 °C and 37 °C, respectively, in either LB or M9 minimal medium (Miller, 1972) with glucose as C-source. Strains were stored in cryotubes at -80 °C, in LB medium containing 15 % glycerol (v/v).

#### Construction of the capacity monitor strain, P. putida CAP

*P. putida* CAP was constructed by inserting the *mCherry* gene into the chromosome of *P. putida* KT2440 under the control of a constitutive promoter. The promoter of the *lac*-operon of *E. coli* was chosen, as this promoter is only regulated by its genomic context (Oehler, 2009). The fragment containing the *mCherry* gene under the control of *lacIp* was cloned into pTn7-M (Zobel et al., 2015) choosing SpeI and SacI restriction sites in *E. coli* DH5α λpir. The resulting pTn7-M_lacIp-*mCherry* vector was introduced into *P. putida* KT2440 by four-parental mating with *E. coli* DH5α λpir (pTn7-M_lacIp-*mCherry*) as donor, *E. coli* HB101 (pRK600) as a helper strain for conjugation, *E. coli* DH5*α* (pTnS1) to provide transposase, and *P. putida* KT2440 as recipient (Choi and Schweizer, 2006; Zobel et al., 2015). The resulting *P. putida* KT2440 *attTn7::lacIp-mCherry* strain, referred to as *P. putida* CAP (for capacity), was selected by growth on citrate and resistance towards gentamycin. The correct integration of *lacIp-mCherry* was confirmed by amplification of a fragment spanning the *lacIp*-*mCherry* region and sequencing.

#### Construction of the burden plasmid pSEVA438-MBPeGFP

The *malE* gene was genetically fused *via* a GlySer-linker (GGGGS) to the N-terminus of *eGFP* by overlap extension PCR. Primers were designed to remove the periplasmic signal sequence of *malE* to avoid extracellular transport of the heterologous protein. eGFP was tagged on the C-terminus with a polyhistidine-tag (6xHis-tag), yielding the MBPeGFP fragment. The fragment was ligated into pSEVA438 using EcoRI and SpeI as restriction sites and the resulting plasmid, pSEVA438-MBPeGFP, was transferred to *E. coli* DH5α λpir using the TSS method (Chung et al., 1989). Positive clones were then selected by streptomycin resistance. After verification of pSEVA438_MBPeGFP by sequencing, the plasmid was transferred to *P. putida* CAP by triparental mating (de Lorenzo and Timmis, 1994), yielding *P. putida* CAP (pSEVA438_MBPeGFP) (Figure 2A).

**Fig. 2.**
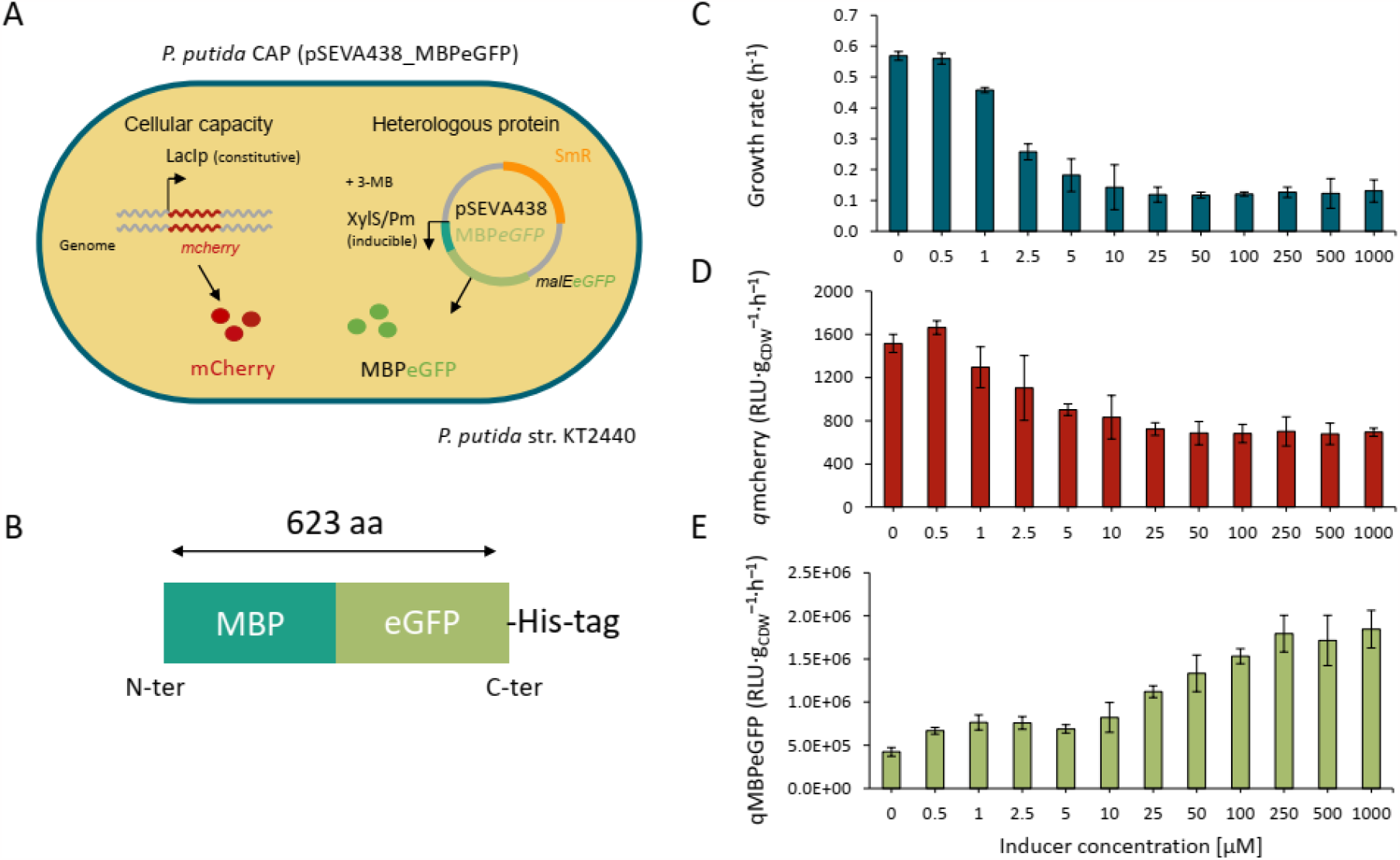
Physiological response of *P. putida* CAP (pSEVA438_MBPeGFP) to heterologous protein production. Schematic representation of the engineered *P. putida* strain with a red fluorescent protein gene (*mCherry*) integrated into the chromosome to monitor biosynthetic capacity and the green fluorescent fusion protein MBPeGFP encoded on a pSEVA438_*MBPeGFP* plasmid (A). Schematic representation of the MBPeGFP fusion protein with an N-terminal maltose binding protein (MBP) domain, the eGFP domain and a C-terminal His-tag sequence (B). Growth rate (C), mCherry production rate (D) and MBPeGFP production rate (E) of *P. putida* CAP (pSEVA438_MBPeGFP) cultivated with different inducer concentrations. Results are averages from at least 3 biological replicates. 3-MB: 3-methyl-benzoate.

### 2.2 Media and preculture conditions

For each experiment, the strains were first streaked onto LB agar plates containing 200 μg/mL streptomycin and incubated overnight at 30 °C. Then, 3 mL of LB medium with 200 μg/mL streptomycin were inoculated from a single isolated colony and the cultures were incubated for 8 to 16 h at 30 °C and 200 rpm in an orbital shaker (Inova 4230, New Brunswick Scientific, New Brunswick, NJ, USA). Cells were first diluted (1/100) in 50 mL of M9 medium containing 17.4 g ·L^−1^ Na_2_HPO_4_ · 12H_2_O, 3.0 g·L^−1^ KH_2_PO_4_, 2.0 g.·L^−1^ NH_4_Cl, 0.5 g.L^-1^ NaCl, 0.5 g·L^−1^ MgSO_4_, and 3.3 mg·L^−1^ CaCl_2_, and 1 mL of a trace element solution containing 15 mg·L^−1^ Na_2_EDTA · 2H_2_O, 4.5 mg·L^−1^ ZnSO_4_ · 7H_2_O, 0.3 mg·L^−1^ CoCl_2_ · 6H_2_O, 1 mg·L^−1^ MnCl_2_ · 4H_2_O, 1 mg·L^−1^ H_3_BO_3_, 0.4 mg·L^−1^ Na_2_MoO · 2H_2_O, 3 mg·L^−1^ FeSO_4_ · 7H_2_O, 0.3 mg·L^−1^ CuSO_4_ · 5H_2_O. M9 preculture medium was supplemented with 3 g·L^-1^ glucose for flux analysis, or with 3 g·L^−1^ of a glucose mixture containing 80 % ^13^C_1_-labeled glucose and 20 % U-^13^C_6_-labeled glucose (Eurisotop, Saint Aubin, France). The glucose, MgSO_4_ and trace element solutions were sterilized by filtration (Minisart 0.2-*μ*m syringe filter; Sartorius, Göttingen, Germany) and other solutions were autoclaved. Cells were incubated in a 250 mL baffled flask at 30 °C and shaken at 200 rpm. Exponentially growing cells were harvested by centrifugation (Sigma 3-18K centrifuge, Sigma-Aldrich, Seelze, Germany) at 5,000 *g* for 15 min at room temperature, washed twice in fresh medium without glucose, and this inoculum was used to inoculate microtiter plates or bioreactors. A 0.5 M stock solution of inducer was prepared by solubilizing 3-methyl-benzoate (3-MB) (Sigma-Aldrich, S^t^ Louis, MO, USA) in 0.5 M NaOH.

### 2.3 Batch culture in microtiter plate

*P. putida* KT2440 CAP (pSEVA438_MBPeGFP) inoculum was diluted to an optical density at 600 nm (OD_600_) of 0.07 (Genesys 6 spectrophotometer, Thermo, Carlsbad, CA, USA) in 5 mL M9 medium containing 3 g·L^-1^ glucose and 200 μg/mL streptomycin and supplemented with 50 *μ*L of different inducer concentrations (0, 0.5, 1, 2.5, 5, 10, 25, 50, 100, 250, 500, 1000 *μ*M) prepared from a 0.5 M stock solution of 3-MB (Sigma-Aldrich, S^t^ Louis, MO, USA) solubilized in a 0.5 M NaOH solution and filtered through a 0.2 *μ*m filter for sterilization (Minisart 0.2 *μ*m syringe filter; Sartorius, Göttingen, Germany). These dilutions were then inoculated (100 *μ*L) in triplicate into a 96 well microtiter plate (Sarstedt, Nümbrecht, Germany) and incubated at 30 °C in a plate reader (CLARIOstar^Plus^, BMG Labtech, Ortenberg, Germany). The optical density at 600 nm (OD_600_) and fluorescence of GFP (excitation wavelength, λex, 470 nm; emission wavelength 1, λem1, 515 nm), autofluorescence (λex, 470 nm; λem, 580 nm) and *mCherry* (λex, 570 nm; λem, 620 nm) were measured every 20 min for 48 h under continuous 200 rpm double orbital shaking. Three independent biological replicates were analyzed.

### 2.4 Batch culture in bioreactors

*P. putida* KT2440 WT and *P. putida* CAP (pSEVA438_MBPeGFP) were grown in a 500 mL bioreactor (my-Control, Applikon Biotechnology INC, Sunnyvale, CA, USA) filled with 300 mL of M9 medium containing 4.0 g·L^−1^ NH_4_Cl, 2.0 g·L^−1^ KH_2_PO_4_, 0.5 g·L^−1^ NaCl, 0.5 g·L^−1^ MgSO_4_, 3.3 mg·L^−1^ CaCl_2_, and 1 mL of the trace element solution supplemented with 10 g·L^−1^ glucose. For flux analysis, ^12^C-glucose was replaced by a 10 g·L^−1^ mixture of 80 % (mol/mol) ^13^C_1_-labeled glucose and 20 % (mol/mol) U-^13^C_6_-labeled glucose (with a ^13^C purity of 99 %; Eurisotop, Saint Aubin, France). The pH was maintained at 7.0 ± 0.1 by automatically adding 14% (g/g) ammonia (VWR, Fontenay-sous-Bois, France) or 10 % (g/g) phosphoric acid (PanReac AppliChem, Barcelona, Spain) and the temperature was set to 30 °C. Adequate aeration of the culture was achieved by automatically controlling the stirrer speed and the gas flow to maintain > 30 % oxygen saturation. Streptomycin (200 *μ*g/mL) was added to the *P. putida* CAP (pSEVA438_MBPeGFP) culture. The inducer for protein production was added before cell inoculation, at 10 *μ*M or 1 mM from a 0.5 M stock solution of 3-MB (Sigma-Aldrich, S^t^ Louis, MO, USA) solubilized in 0.5 M NaOH. Foaming was avoided by manually adding 50 *μ*L of antifoam 204 (Sigma-Aldrich, S^t^ Louis, MO, USA) before inoculation and when required during growth. The bioreactor was inoculated to an OD_600_ of 0.15. Depending on the culture time, 1.0 to 2.5 mL of culture were withdrawn from the bioreactor every 0.5 to 1 h to monitor the OD_600_, fluorescence, and protein and metabolite contents of the culture. Cell dry weight (CDW) was determined by filtering 5 mL of cells grown in M9 supplemented with glucose at OD_600_ 1.0, 1.5 and 2.0 on preweighed membrane filters (Sartorius 0.2 *μ*m, 47 mm, Goettingen, Germany) under vacuum filtration. The filters were then washed with a 0.9 % NaCl solution, incubated at 80 °C and weighed every 24 h until a constant weight was reached. Under our laboratory conditions, the OD_600_ CDW correlation for *P. putida* was CDW [g·L^−1^] = 0.57 ± 0.01 × OD_600_.

### 2.5 Fluorescence data processing

The green fluorescence specifically emitted by eGFP was measured in relative light units (RLU_GFP_) using a procedure similar to the one described by Lichten et al. (2014). The autofluorescence signal emitted by growing cells was estimated by determining the ratio (*r*_a_) of fluorescence at λem = 515 nm – specific to eGFP fluorescence signals – and 580 nm (non-specific to eGFP fluorescence signals) of a *P. putida* KT2440 WT culture grown in microtiter plate, yielding a *r*_a_=RLU_515_/RLU_580_ = 2.479 (R^2^=0.999; Figure S1A supplementary data). Next, based on these data, the fluorescence emitted at 515 nm and at 580 nm by a culture of *P. putida* CAP (pSEVA438_MBPeGFP) was measured at each time point and the autofluorescence was subtracted from the level of green fluorescence, yielding the amount of fluorescence specific to the heterologous protein, RLU_GFP_= *RLU*_515_-*r*_a_x*RLU*_580_, (Lichten et al., 2014).The overlap between the autofluorescence signal of the cells and the fluorescence emitted by mCherry at 620 nm was negligible.

### 2.6 Purification of heterologous protein

The heterologous protein MBPeGFP was purified using NEBExpress Ni spin columns (NEB, Ipswich, MA, USA) according to manufacturer recommendations. Briefly, 50 mL of *P. putida* CAP (pSEVA438_MBPeGFP) culture induced with 1 mM of 3-MB (Sigma-Aldrich, S^t^ Louis, MO, USA) were extracted from a bioreactor at OD_600_ of 2.25, harvested at 5,000 *g* for 15 min, and stored at -80 °C until further analysis. Frozen cell pellets were resuspended in 4 mL of IMAC buffer (NEB, Ipswich, MA, USA) on ice and dispersed using an ultrasonic disruptor (Sonics, VibraCell™, Newtown, CT, USA) in 9 sonication cycles (20 s ON, 30 s OFF; power amplitude, 20 %) with a pause of 60 s in between. Cell debris was removed by centrifugation (15 min at 16,100 *g*). Protein purification was then carried out following the manufacturer’s protocol. After adding 200 *μ*L of elution buffer containing 50 mM imidazole, the concentration of MBPeGFP was measured with a bicinchoninic acid (BCA) assay using bovine serum albumin (BSA) dilutions ranging from 0 to 206.25 *μ*g/mL to establish a standard curve (Figure S1B). Purified MBPeGFP was two-fold serially diluted. BSA standard and purified protein (25 *μ*L) were each added in triplicate to a microtiter plate (Sarstedt, Nümbrecht, Germany) and 200 *μ*L of a 50:1 mixture of BCA and copper(II) sulfate solutions (Sigma-Aldrich, S^t^ Louis, MO, USA) was added to each well containing a sample. The plate was incubated for 30 min at 37 °C, then left to cool at room temperature for 5 min before absorbance was measured at 562 nm with a plate reader (CLARIOstar^Plus^, BMG Labtech, Ortenberg, Germany). The final concentration of purified protein was 1076 *μ*g/mL. This stock solution was then diluted (at concentrations of 175, 150, 125, 100, 75, 50 *μ*g/mL) and mixed with 4X Laemmli buffer (Biorad, Hercules, CA, USA) containing 1 % β-mercaptoethanol for SDS-PAGE calibration, as detailed below (Figure S1C and S1D).

### 2.7 Heterologous protein quantification by SDS-PAGE and fluorescence correlation factor

Bioreactor culture samples (60 *μ*L) were extracted at an OD_600_ of 1 to 4, diluted to an OD_600_ of 1, mixed with 4X Laemmli buffer (Biorad, Hercules, CA, USA) containing 1 % β-mercaptoethanol and stored at −20 °C until analysis. After denaturation for 5 min at 95 °C, 8 μL of cellular extract and dilutions of purified protein (at 175, 150, 125, 100, 75, 50 *μ*g/mL) were loaded onto a 12.5 % SDS-PAGE gel and separated at 160 V for 70 min in a mini-protean tetra cell (Biorad, Hercules, CA, USA) using Tris-glycine-SDS running buffer. The gels were washed once with distilled water and stained with InstantBlue® Coomassie Protein Stain (Abcam, Cambridge, UK). The gels were viewed using a ChemiDOC XRS molecular imager (Biorad, Hercules, CA, USA) (Figure S1C) and the images were analyzed using Image Lab v6.0.1 (Biorad, Hercules, CA, USA). Brightness and contrast were adjusted automatically using the software. Lanes and bands were identified manually. The lane profile tool was used to quantify the peak area that corresponded as precisely as possible to the protein band. The resulting calibration curve for MBPeGFP (Figure S1D) was used to estimate its concentration in the samples (Figure S1E). A correlation factor between the amount of green fluorescence (RLU_MBPeGFP_) emitted by the sample and the protein concentration was calculated from the slope of the curve (Fluo factor 7.25 x 10^-9^ - 0.0007, R^2^ = 0.954). This was used to convert the levels of green fluorescence measured in other samples into MBPeGFP protein concentrations.

### 2.8 Quantification of extracellular metabolites by NMR

Glucose, gluconate and 2-KG were identified and quantified by nuclear magnetic resonance (NMR). Culture samples (1 mL) were collected every 0.5 to 1 h during growth, centrifuged at 14,500 *g* for 3 min, and the supernatants were stored at −20 °C until analysis. The supernatants (180 *μ*L) were mixed with 20 *μ*L of an internal standard consisting of 10 mM deuterated trimethylsilylpropanoic acid (TSP-d4) diluted in D_2_O. Proton NMR spectra were recorded on an Avance III 500-MHz spectrometer equipped with a 5-mm z-gradient TXI (^1^H, ^13^C, ^31^P) probe (Brucker, Rheinstatten, Germany). Quantitative ^1^H NMR analysis was performed at 286 K, using a zgpr30 sequence with a recycle delay of 10 s, and presaturation at 4.7 ppm for water suppression. Thirty-two scans were accumulated (32k data points with a spectral width of 10 ppm) after 4 dummy scans. Inverse gated ^13^C decoupling was used for samples containing ^13^C-Glucose to ensure the spectra were quantitative. The zgigpr sequence was used with ^13^C decoupling during ^1^H acquisition (396 ms), a recycle delay of 10 s, and presaturation at 4.7 ppm for water suppression. Thirty-two scans were accumulated (8k data points with a spectral width of 20 ppm). The spectra were processed using Topspin 3.1 (Bruker, Rheinstatten, Germany). The reported results are the average values obtained from at least three biological replicates.

### 2.9 Calculation of Growth Rates and Extracellular Fluxes

Growth rates and extracellular metabolite fluxes were determined in the exponential growth phase from the time courses of biomass, glucose, gluconate and 2-KG concentrations, as measured by NMR. All calculations were performed using PhysioFit 1.0.2 (Peiro et al., 2019) (https://github.com/MetaSys-LISBP/PhysioFit) from culture profiles obtained with unlabeled or ^13^C-labeled glucose (Figure S2). MBPeGFP and mCherry production rates (*q*_MBPeGFP_ and *q*_mcherry_) were calculated by multiplying the respective yields by the growth rate.

### 2.10 ^13^C-labeled samples and quantitative isotopic analysis

Samples of culture medium (100 *μ*L) for quantitative isotopic analyses and ^13^C-metabolic flux analyses were collected during the exponential growth phase (between OD_600_ 2 and 4) from culture grown on ^13^C-glucose. The samples were plunged and vigorously mixed in 2 mL of methanol/acetonitrile/H_2_O (4:4:2) precooled at -20 °C, incubated for 2 h at -20 °C, evaporated overnight in a SpeedVac (SC110A SpeedVac Plus, ThermoSavant, Waltham, MA, USA) and stored at -80 °C until IC-MS analysis. The cell extracts were then resuspended in 100 *μ*L deionized water, and centrifuged at 14,500 *g* for 10 min at 4 °C to remove cell debris. Mass fractions of intracellular metabolites were quantified using an ion chromatograph (IC; Thermo Scientific Dionex ICS-50001 system; Dionex, Sunnyvale, CA, United States) coupled to an LTQ Orbitrap mass spectrometer (Thermo Fisher Scientific, Waltham, MA, United States) equipped with a heated electrospray ionization (HESI) probe. Fourier transform mass spectra were recorded in full-scan negative ion mode at a resolution of 60,000 at m/z = 400. The ion chromatography and mass spectrometry experiments are described in greater detail elsewhere (Vogeleer and Létisse, 2022). In total, three samples were analyzed for each of three independent biological replicates.

Isotopologues were quantified from 11 metabolic intermediates covering the central metabolism (6-phosphogluconate, glucose 6-phosphate, fructose 6-phosphate, fructose 1,6-bisphosphate, 2/3-phosphoglycerate, phosphoenolpyruvate, ribose 5-phosphate, sedoheptulose 7-phosphate, citrate/isocitrate, malate and succinate). The mass fractions were corrected for naturally occurring isotopes, using IsoCor 2.2.0 ((Millard et al., 2019), https://github.com/MetaSys-LISBP/IsoCor) and high resolution spectra (60,000 at m/z = 400), with corrections for the natural abundance and isotopic purity (99 %) of the tracer element.

### 2.11 ^13^C-Metabolic flux analyses

^13^C metabolic fluxes were calculated using the software influx_s (Sokol et al., 2012), based on a flux model that included the mass balances and carbon atom transitions of *P. putida*’s main central metabolic pathways (Kohlstedt and Wittmann, 2019; Nikel et al., 2015): glucose uptake, glycolysis (EMP), pentose phosphate pathway (PPP), ED pathway, tricarboxylic acid cycle (TCA), anaplerotic reactions, and gluconate and 2-KG secretion. Following Kohlstedt and Wittmann (2019), the flux through the Kdgk reaction was fixed at 4.9 % of the flux through the Gntk reaction. Precursor requirements for biomass synthesis were estimated based on the composition of the biomass (van Duuren et al., 2013) and growth rates. Similarly, precursor requirements for the synthesis of heterologous protein were estimated from the protein sequence and from protein production rates. Intracellular fluxes were estimated by fitting i) extracellular fluxes (glucose uptake, and gluconate, 2-KG, biomass and protein production) and ii) the ^13^C-labelling patterns of intracellular metabolites measured by IC-MS. A chi-square statistical test was used to assess the goodness-of-fit (based on 95 % confidence intervals) for each condition. For comparison, the estimated fluxes were then normalized to the rate of substrate uptake, set to 100. For each condition, the mean fluxes and the corresponding standard deviations were determined from three independent biological replicates. The isotopic data and metabolic fluxes for each independent biological replicate are given in Table S2 in the supplementary material. All model files and flux calculation results are available at https://github.com/MetaSys-LISBP/pseudomonas_metabolic_burden.

### 2.12 Calculation of carbon, redox and energy balances

The carbon balance was determined by calculating the proportion of carbon converted from glucose to biomass, gluconate, 2-KG, CO_2_ and heterologous protein during the exponential phase as outlined in the Results section. The amounts of biomass, gluconate, 2-KG and heterologous protein produced were quantified as described above. The amount of CO_2_ produced during the exponential phase was calculated from the percentage concentrations of CO_2_ and N_2_ measured in the gas output using a Dycor ProLine Process mass spectrometer (Ametek, Berwyn, PA, USA). The biomass formula was assumed to be C_1_H_1.52_O_0.41_N_0.27_P_0.02_S_0.01_ (van Duuren et al., 2013).

NADPH production and consumption fluxes were estimated by summing the fluxes through all reactions producing or consuming NADPH along with NADPH fluxes for the synthesis of biomass (anabolism) and heterologous protein. The apparent excess of NADPH may be converted to NADH through the activities of pyridine nucleotide transhydrogenases (SthA and PntAB) (Nikel et al., 2016b), as shown in previous work (Kohlstedt and Wittmann, 2019). Similarly, ATP production *via* substrate-level phosphorylation was calculated by summing the fluxes of ATP-producing reactions and subtracting the fluxes of ATP-consuming reactions. ATP produced *via* oxidative phosphorylation was inferred from the production rates of NADH, FADH_2_ and FMNH_2_, assuming P/O ratios of of 1.875 for the conversion of NADH and PQQH_2_ into ATP and a P/O ratio of 1.0 for the conversion of FADH_2_ (Nikel et al., 2015). ATP demand was calculated by summing the requirements for anabolism, non-growth-associated maintenance (van Duuren et al., 2013), and for the biosynthesis of the heterologous protein, assuming 5 ATP equivalents per amino acid residue added (Garett and Grisham, 1999).

## 3. Results

### 3.1 Characterization of the metabolic burden in *P. putida*

To characterize the physiological impact of the metabolic burden due to heterologous protein production in *P. putida* KT2440, we introduced a fluorescent protein under the control of a constitutive promoter into the *att* site in the chromosome as a proxy for the biosynthetic capacity of *P. putida* cells. We chose the mCherry fluorescent protein controlled by the *lacIp* promotor from the *E. coli lac*-operon, which is not regulated when isolated from its original genomic environment. The amount of mCherry produced during heterologous protein production was taken to be representative of the availability of cellular resources and served as a proxy for the biosynthetic capacity of the cells (Ceroni et al., 2015). The reasoning behind is that the amount of fluorescent protein produced from the non-regulated promoter should resemble the amount of protein produced from the native constitutive promoters in the genome. Thus, it can be understood as a stand-in for the availability of cellular resources, such as RNAP, Ribosomes, tRNAs, or amino acids.

The eGFP labeled protein chosen to impose a metabolic burden was introduced into *P. putida* KT2440 cells on a plasmid under the control of an inducible promoter (Figure 2A). The eGFP domain was fused to the C-terminus of MBP, forming a 623-amino-acid fusion protein, MBPeGFP (Figure 2B). The advantages of MBPeGFP for this study were that the fluorescence emitted by eGFP could be used to estimate protein yields during growth, while the MBP domain increased solubility and limited aggregation (Raran-Kurussi et al., 2015). To facilitate purification, a His-tag was added to the C-terminal part of the eGFP domain (sequence is available in supplementary data). The MBPeGFP encoding gene was integrated into pSEVA438, under the control of a XylS/Pm expression system induced by 3-MB.

The associated metabolic burden was quantified in terms of the growth, biosynthetic capacity and heterologous protein production of *P. putida* CAP (pSEVA438_MBPeGFP) cultures in microtiter plates. These small-scale experiments demonstrated that both the mCherry production and the maintenance of the pSEVA438 plasmid did not impose a noticeable metabolic burden (Figure S3). We rather assumed that these processes consume part of the metabolic leeway. Next, the production of MBPeGFP was modulated by exposing the cells to various concentrations of 3-MB inducer, from 0 to 1000 *μ*M.

Four types of behavior can be distinguished in these data. First, at very low 3-MB concentrations (< 0.5 *μ*M), the small amounts of MBPeGFP produced did not represent a metabolic burden since there was no impact on *P. putida* growth or biosynthetic capacity (mCherry production). However, some MBPeGFP production was observed even in the absence of inducer, indicating that as reported previously (Balzer et al., 2013), the XylS/Pm expression system is slightly leaky. The growth rate (*μ*_max_) was 0.57 ± 0.02 h^-1^ (Figure 2C), as reported in the literature for this strain when grown on glucose (Kohlstedt and Wittmann, 2019; Kozaeva et al., 2021). Second, exposure to between 0.5 and 10 *μ*M 3-MB was associated with a dramatic reduction in *μ*_max_ from 0.57 ± 0.02 h^-1^ to 0.15 ± 0.02 h^-1^, and decreased mCherry production (Figure 2D). This relationship between growth rate and biosynthetic capacity is in agreement with Ceroni et al.’s findings in *E. coli* (2015) . The MBPeGFP production rate remained constant, meaning that the decrease in cell growth did not directly translate into a higher heterologous protein production rate (Figure 2E). Under these conditions, resource sharing is detrimental to cell growth with no clear benefit in terms of protein production. From 10 to 250 *μ*M 3-MB, the MBPeGFP production rate gradually increased while cell growth and mCherry production remained constant. The MBPeGFP production rate at 250 *μ*M 3-MB was twice what it was at 10 *μ*M 3-MB. Here too, the rates of cell growth and protein production did not directly counterbalance each other, indicating that the coordination of these processes is more complex than a simple sharing of cellular resources. Finally, exposing the cells to more than 250 *μ*M 3-MB did not increase the MBPeGFP production rate and had no additional impact on their growth rate or biosynthetic capacity.

Having observed that exposing *P. putida* CAP (pSEVA438_MBPeGFP) to various concentrations of inducer generated different levels of metabolic burden, we explored how the central metabolism of the host copes with *low* (without inducer), *medium* (10 *μ*M 3-MB), and *high* (1000 *μ*M 3-MB) degrees of metabolic burden. We chose the *P. putida* KT2440 wild-type strain as a reference in this study, as there was no major difference in either growth rate or capacity - where applicable - between WT and CAP strains harboring or not the empty plasmid (Figure S3). On the other hand, quantitative data on metabolic fluxes are available for this strain in the literature, which allows us to compare and validate our methodology for analyzing metabolic fluxes (Kohlstedt and Wittmann, 2019).

### 3.2 Quantitative analysis of *P. putida* physiology during heterologous protein production

Cell cultures were scaled-up from micro-titer plates to bioreactors so that key environmental parameters (pH, dissolved oxygen tension (DOT), temperature, agitation, *etc*.) could be adjusted precisely. *P. putida* CAP (pSEVA438_MBPeGFP) was cultivated in minimal medium supplemented with 10 g·L ^-1^ glucose as sole carbon source and containing either none, 10 *μ*M or 1000 *μ*M 3-MB. Cells were grown in the presence of unlabeled or ^13^C-labeled glucose (Figure S2) and growth parameters and carbon balances (Table 1 and Figure 3) were determined from the growth profiles. Intracellular fluxes were determined from the cultures supplemented with ^13^C-labeled glucose (see paragraph 3.5).

**Table 1:** Growth parameters of glucose-grown *P. putida* KT2440 WT and *P. putida* CAP (pS EVA438_MBPeGFP) for different inducer concentrations.

| Parameter | WT strain<br>(n=3)<sup>b</sup> | *P. putida* CAP (pSEVA438-MBPeGFP)<br>(n=4)<sup>b</sup> |  |  |
| --- | --- | --- | --- | --- |
| 3-methyl-benzoate (μM) | 0 | 0 | 10 μM | 1000 μM |
| $\mu_{\max}$ [(h<sup>-1</sup>)] | 0.58 ± 0.02 | 0.58 ± 0.03 | 0.29 ± 0.02 | 0.19 ± 0.04 |
| $q_{\text{mcherry}}$ (RLU · [g<sub>CDW</sub> · h]<sup>-1</sup>) | ND<sup>a</sup> | 1150 ± 101 | 928 ± 48 | 555 ± 112 |
| $q_{\text{MBPeGFP}}$ (mg · [g<sub>CDW</sub> · h]<sup>-1</sup>) | ND<sup>a</sup> | 2.8 ± 0.5 | 9.9 ± 2.9 | 26.2 ± 8.5 |
| $q_{\text{CO}_2}$ (mmol · [g<sub>CDW</sub> · h]<sup>-1</sup>) | 13.0 ± 1.9 | 12.5 ± 1.9 | 10.8 ± 1.3 | 9.0 ± 1.3 |
| $Y_{X/S}$ (g<sub>X</sub> · mol<sup>-1</sup>) | 82.1 ± 6.8 | 84.4 ± 4.2 | 68.7 ± 3.9 | 55.0 ± 7.0 |
| $Y_{\text{MBPeGFP}}$ (mg · g<sub>CDW</sub><sup>-1</sup>) | ND<sup>a</sup> | 4.7 ± 1.0 | 34.9 ± 11.4 | 136.2 ± 19.8 |
| $q_{\text{Glc}}$ (mmol · [g<sub>CDW</sub> · h]<sup>-1</sup>) | 7.0 ± 0.3 | 6.9 ± 0.5 | 4.2 ± 0.3 | 3.4 ± 0.3 |
| $q_{\text{Gnt}}$ (mmol · [g<sub>CDW</sub> · h]<sup>-1</sup>) | 0.78 ± 0.22 | 0.81 ± 0.18 | 0.37 ± 0.16 | 0.15 ± 0.07 |
| $q_{2\text{-KG}}$ (mmol · [g<sub>CDW</sub> · h]<sup>-1</sup>) | 0.06 ± 0.01 | 0.02 ± 0.02 | 0.21 ± 0.03 | 0.22 ± 0.02 |

<sup>a</sup> Not detected
<sup>b</sup> The average values and standard errors of the means were calculated from the values measured in *n* biological replicates.

**Fig. 3:**
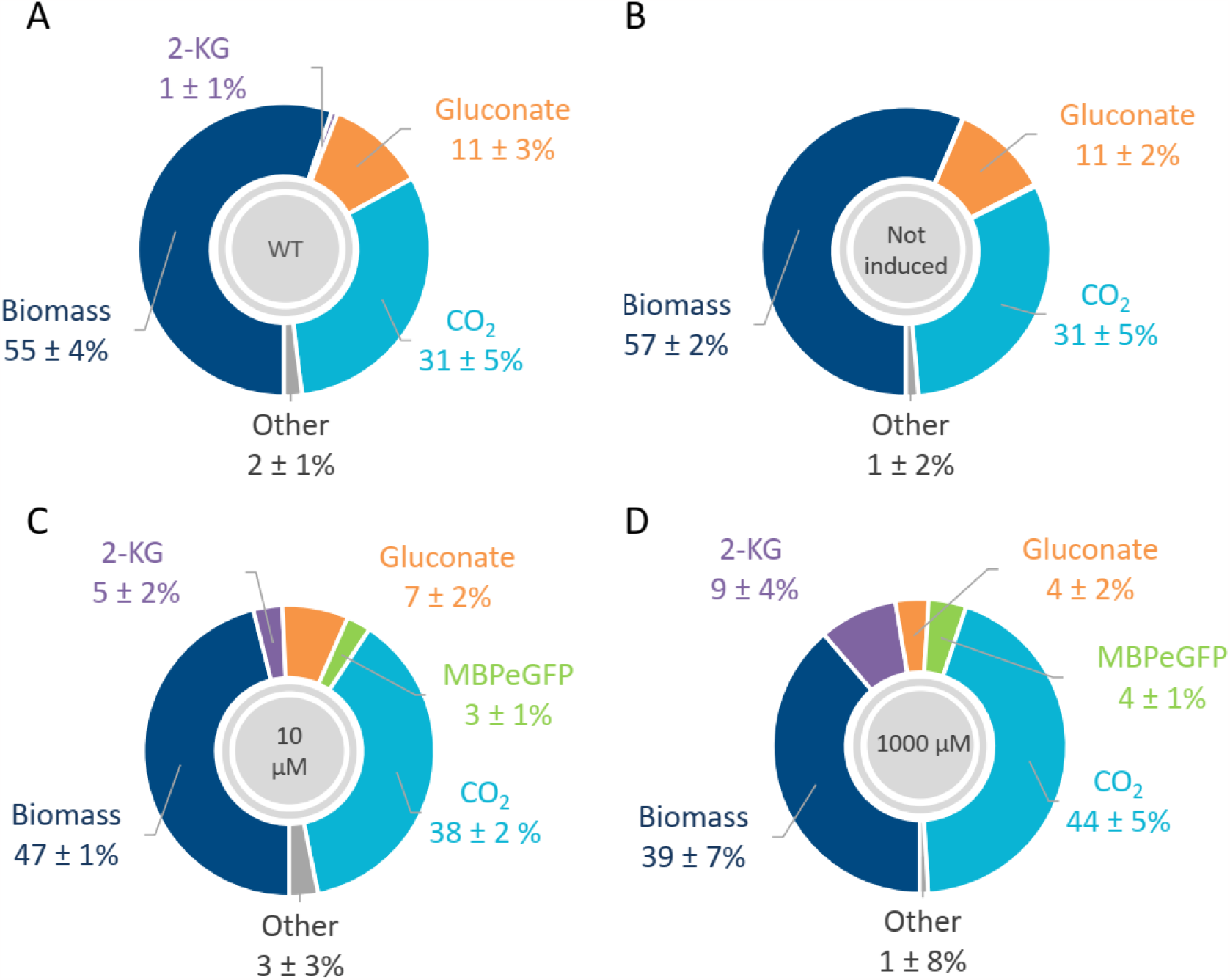
Carbon balance of glucose-grown *P. putida* KT2440 WT (A) and *P. putida* CAP (pSEVA438_MBPeGFP) under low (B), medium (C) and high (D) heterologous protein production conditions. Carbon balances were calculated from the concentrations measured in the media over exponential growth phase, highlighted in blue in Figure S2. Values represent the mean ± standard deviation of three biological replicates for WT strain and four biological replicates for CAP (pSEVA438_MBPeGFP) strain.

In the absence of inducer, *P. putida* CAP (pSEVA438_MBPeGFP) grew at the same rate (0.58 ± 0.02 h^-1^) in the bioreactor as in the microtiter plate (Figure 2) and as the WT strain in the bioreactor (Table1). This value is also similar to those reported in the literature for growth on glucose (Kohlstedt and Wittmann, 2019). In the presence of 10 or 1000 *μ*M 3-MB, the growth rates in the exponential phase were half (0.29 ± 0.02 h^-1^) and one third (0.19 ± 0.04 h^-1^) of the non-induced value, respectively. As observed in microtiter plates, the production rate of mCherry also decreased, by 20% and 50%, respectively.

Under non-induced conditions, MBPeGFP represented an estimated 0.47± 0.1 % of the total CDW, confirming the slight leakiness of the XylS/Pm expression system. In the presence of 10 *μ*M and 1000 *μ*M inducer, the production rate of MBPeGFP increased by factors of almost 4 and 10, respectively, leading to heterologous protein yields of 3.49 ± 1.1 % and 13.6 ± 2.0 % of the total CDW (Table 1).

These results show that the chosen induction conditions had the expected effects, namely a *low* metabolic burden in the absence of induction, and *medium* and *high* burdens with 10 and 1000 *μ*M 3-MB, respectively.

### 3.3 Protein production modifies the periplasmic metabolism

The burden imposed by the synthetic expression system for maintenance and protein production leads to profound modifications of the central metabolism (Wittmann et al., 2007). To investigate the metabolic rearrangements occurring in *P. putida* in response to heterologous protein production, the dynamic profiles of extracellular metabolites in the culture media of WT and CAP (pSEVA438_MBPeGFP) *P. putida* strains were quantified by NMR.

*P. putida* has been observed to produce gluconate and small amounts of 2-KG during exponential growth on glucose, with concentrations peaking at 5 mM and 0.2 mM, respectively (Figure S2), and both compounds subsequently being co-consumed with glucose (Nikel et al., 2015). In the absence of inducer, a similar profile was observed here for gluconate, while the 2-KG concentration dropped and approached the limit of detection (Figure S2). The concentrations of both compounds reached much higher levels in the presence of inducer, with the concentration of 2-KG peaking at 2 and 4 mM in the presence of 10 and 1000 *μ*M 3-MB, respectively. No other metabolites were found to accumulate in these experiments.

The rates of glucose consumption (q_Glc_), gluconate and 2-KG production (q_Gnt_ and q_2KG_), and mCherry and MBPeGFP production (q_mcherry_, q_GFP_) were determined during the exponential growth phase based on the concentrations of biomass and extracellular metabolites and on protein fluorescence. The fact that these rates were constant throughout the exponential growth phase indicates that the cells had reached a steady state (Table 1 and Figure S4). Under low MBPeGFP production conditions, the rates of glucose consumption (6.9 mmol·g^−1^·h^−1^ ± 0.5) and gluconate production (0.81 ± 0.18 mmol·g^−1^·h^−1^) were similar to those calculated for the WT strain (Table 1) (Kohlstedt and Wittmann, 2019). Under medium and high MBPeGFP production conditions, glucose consumption decreased to 4.2 ± 0.3 mmol·g^−1^·h^−1^ in the presence of 10 *μ*M 3-MB and 3.4 ± 0.3 mmol·g^−1^·h^−1^ at 1000 *μ*M 3-MB, while 2-KG production was close to 0.2 mmol·g^−1^·h^−1^ under both conditions (Table 1). CO_2_ production was also lower under these conditions than in the WT strain.

These results indicate that heterologous protein production caused a marked slowdown of metabolic activity in *P. putida* CAP (pSEVA438_MBPeGFP), as reflected in particular by the lower rate of glucose consumption. More surprising is the sharp change in periplasmic metabolism evidenced by the accumulation of 2-KG. To the best of our knowledge, this phenomenon has never previously been reported in *P. putid*a, but it is similar to the acetate overaccumulation observed under recombinant protein production conditions in *E. coli* (San et al., 1994; Ying Lin and Neubauer, 2000).

### 3.4 Heterologous protein production consumes very little carbon but profoundly alters the distribution of carbon usage in *P. putida*

The carbon balance was calculated for each condition by summing the amounts of carbon used to produce biomass, gluconate, 2-KG, CO_2_ and heterologous protein (Figure 3). These compounds accounted for nearly 100 % of carbon usage in each case (WT: 98 ± 1 %, uninduced: 99 ± 2 %, 10 *μ*M: 100 ± 4 %, 1000 *μ*M: 98 ± 8 %), confirming that no other carbon molecules were produced in significant amounts. Under low MBPeGFP production conditions, the proportions of carbon used for biomass, gluconate, CO_2_ and 2-KG production were similar to those observed for the WT strain. The proportion of carbon used for biomass production decreased to 0.46 ± 0.01 Cmol/Cmol and 0.39 ± 0.07 Cmol/Cmol under medium and high protein production conditions, respectively, while the proportion of carbon diverted to gluconate decreased by a factor of 1.6 and 2.8, respectively, and the fraction used for 2-KG production increased significantly to 5 ± 2 % of total carbon usage at 10 *μ*M 3-MB and 9 ± 4 % 1000 *μ*M 3-MB. Remarkably, the total amount of carbon converted into gluconate and 2-KG was constant across all studied conditions.

The proportion of carbon used for MBPeGFP production was negligible in the absence of inducer, and was just 0.03 ± 0.01 Cmol/Cmol and 0.04 ± 0.02 Cmol/Cmol in the presence of 10 and 1000 *μ*M 3-MB, respectively. However, the amount of carbon excreted as CO_2_ by the cell increased from 0.31 ± 0.05 Cmol/Cmol without inducer to 0.38 ± 0.02 Cmol/Cmol in the presence of 10 *μ*M 3-MB and 0.44 ± 0.05 Cmol/Cmol with 1000 *μ*M 3-MB (Figure 3). In other words, under conditions of significant heterologous protein production, part of the carbon normally used for biomass was completely oxidized into CO_2_. Therefore, although production of the heterologous protein only accounted for a small percentage of the total carbon consumed, it had a profound impact on cellular metabolism.

### 3.5 Metabolic flux analyses reveal metabolic rearrangements in the central metabolism of *P. putida*

We employed ^13^C metabolic flux analysis to measure the alterations in the central metabolism of *P. putida* in response to heterologous protein production. To minimize the impact of culture history, we quantified ^13^C incorporation in central metabolites that have a faster turnover rate compared to the end products of the carbon metabolism. Data was collected from the cultures grown in the presence of ^13^C-labeled glucose (Figure S2).

#### 3.5.1 Metabolic flux map for *P. putida* WT

We first compared the central metabolic flux distribution measured in WT *P. putida* KT2440 (Figure 4) with those based on LC- or GC-MS based ^13^C metabolic flux analyses reported in the literature (Nikel et al., 2015; Kohlstedt and Wittmann, 2019). Similarly to previous reports, we found that glucose was mainly oxidized in the periplasm into gluconate (91 %) by glucose dehydrogenase (Gcd) and to a lesser extent (5 %) into 2-KG by gluconate dehydrogenase (Gad), with a significant (> 10 %) gluconate excretion into the culture supernatant. Most of the gluconate (76 %) was phosphorylated into 6-phosphogluconate (6PG). At this metabolic node, the carbon flux through the 2-KG bypass was 4 %. 6PG was mainly channeled into the ED pathway *via* 2-keto-3-deoxy-6-phosphogluconate (2-KDPG), cleaved into glyceraldehyde-3-phosphate (G3P) and pyruvate. G3P was mainly driven toward the low EMP pathway, where it converged with 2-KDPG breakdown at the pyruvate node. Approximatively 20 % of G3P was directed toward the EDEMP cycle. Despite this flux, glucose was mainly oxidated in the periplasm, where the oxidative flux was 6 times higher than in the cytoplasm. The flux channeled through the EDEMP cycle is however essential as it increases the NADPH reducing power and ensures the supply of precursors (R5P, E4P, F6P) required for anabolism. Finally, pyruvate was mainly driven toward the TCA cycle, with no significant flux through the glyoxylate shunt. The concerted action of pyruvate dehydrogenase (Pdh) and pyruvate carboxylase (Pyc) fuels the TCA cycle while replenishing the supply for anabolic needs. The flux through malic enzyme (Mae) was remarkably high, contributing approximately one quarter of the total influx into the pyruvate pool.

**Fig. 4.**
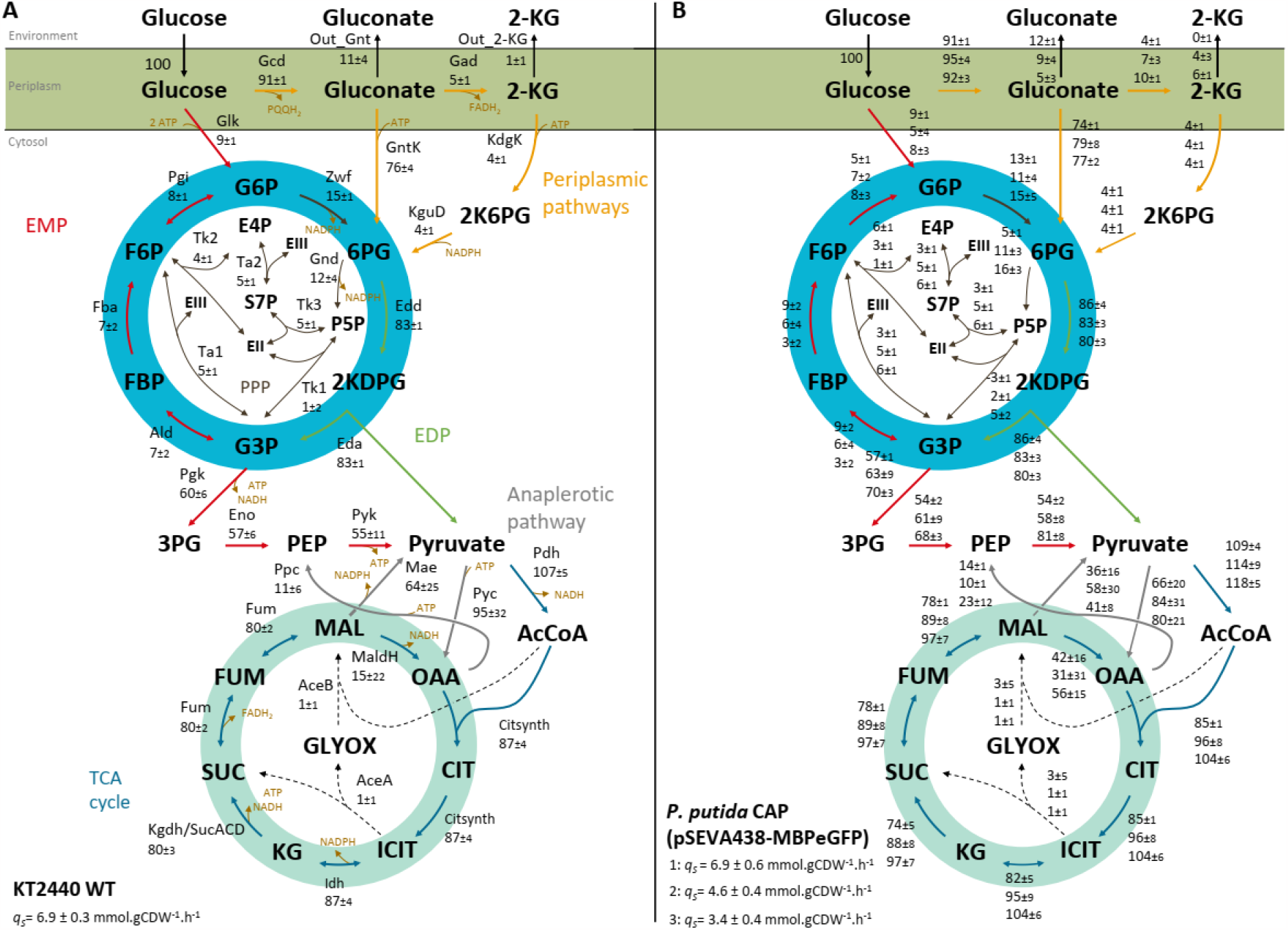
Relative carbon flux distributions of KT2440 WT (A) and *P. putida* CAP (pSEVA438_MBPeGFP) (B) grown on glucose with or without 3-MB. All flux values are normalized to the specific glucose uptake rate of each condition, which was set to 100. Values represent the mean ± standard deviation of two biological replicates for the WT strain (A) and 3 biological replicates for the CAP (pSEVA438_MBPeGFP) strain (B). In panel B, the flux values correspond from top to bottom to: low (without inducer), medium (10 *μ*M 3-MB), and high (1000 *μ*M 3-MB) heterologous protein production conditions. Reaction names and equations are provided in the supplemental data (Table S1).

#### 3.5.2 Metabolic flux map for *P. putida* CAP (pSEVA438_MBPeGFP)

Next, we examined the metabolic flux distributions in the central metabolism of *P. putida* CAP (pSEVA438_MBPeGFP) under different degrees of metabolic burden.

As mentioned above, *P. putida* CAP (pSEVA438_MBPeGFP) produced small amounts of heterologous protein even in the absence of inducer, exploiting only a part of its free capacity. The flux distribution in the absence of inducer was similar to the one determined for the WT strain (Figure 4B), indicating that using a small amount of *P. putida*’s free capacity only has a negligible effect on the carbon flux distribution. In other words, this metabolic leeway can be exploited to produce exogenous compounds such as proteins without affecting the cellular machinery. Increasing this free capacity would therefore appear to be a viable strategy to increase the amount of heterologous protein that can be produced without imposing a metabolic burden. For example, removing 4.3 % of dispensable genes from *P. putida*’s genome (including genes encoding for flagellar motility and genome stability) has been observed to improve heterologous protein yields by 40 % (Lieder et al., 2015).

In contrast, the central metabolic flux distribution of *P. putida* CAP (pSEVA438_MBPeGFP) was profoundly altered under medium and high metabolic burdens. First, periplasmic gluconate oxidation into 2-KG by FAD-dependent Gad increased up to 2-fold without any significant change either in gluconate oxidation through the ED pathway *via* gluconate kinase (GntK) or *via* 2-KG bypass (Figure 4B). The increased flux through Gad was compensated by a 2-fold decrease in excreted gluconate and by significant excretion of 2-KG. Therefore, this metabolic reshuffling resulted mainly in increased production of FADH_2_ through Gad flux.

Second, whereas the flux through the EDEMP cycle tended to decrease under increased MBPeGFP production, the flux through the other pathways of the central metabolism increased, particularly through the endergonic subset of the EMP pathway and in the TCA cycle. The fluxes through phosphoglycerate kinase (Pgk), phosphopyruvate hydratase (Eno) and pyruvate kinase (Pyk) increased by respectively 10 %, 12 % and 7 % under medium MBPeGFP production compared with the low production condition, and by 22 %, 26 % and 50 % respectively at high MBPeGFP production (Figure 4A and 4B). The fluxes through the TCA cycle dehydrogenases (isocitrate dehydrogenase (Idh), *α*-ketoglutarate dehydrogenase (AkgdH), succinate dehydrogenase (Sdh) and to a lesser extent malate dehydrogenase (Mdh)) increased by roughly 15% and 25% under medium and high MBPeGFP production conditions relative to the baseline level. In addition, the glyoxylate cycle was not activated by protein production.

This analysis of metabolic fluxes shows that the production of the heterologous protein MBPeGFP leads to a reshuffling of *P. putida*’s central carbon metabolism, with carbon fluxes redirected toward the oxidative catabolic pathway. This presumably provides the extra chemical and redox energy required to synthesize the heterologous protein and counter the corresponding stress while still satisfying basic housekeeping needs.

### 3.6 *P. putida* generates an apparent excess of ATP under heterologous protein production conditions

The rearrangement of central carbon fluxes for heterologous protein production point to an important adjustment of energy metabolism. We therefore calculated redox and ATP fluxes from the measured flux distributions as described in the Materials and Methods section.

In agreement with previous studies (Kohlstedt and Wittmann, 2019; Nikel et al., 2015), our data indicate that *P. putida* WT generates a catabolic excess of NADPH (Figure 5A). Isocitrate dehydrogenase (Idh) supplies roughly half of the NADPH, and malic enzyme a third. The oxidative branch of the PP pathway contributes very little, despite it being the main contributor to NADPH production in other microorganisms, such as *E. coli* (Nicolas et al., 2007). The apparent surplus of NADPH is converted into NADH *via* transhydrogenase activities (PntAB and SthA). As expected in aerobic growth, ATP is overwhelmingly produced by oxidative phosphorylation, mainly for anabolic purposes (Figure 5B). Unlike NADPH, ATP-production and consumption balance out (>93 %).

**Fig. 5.**
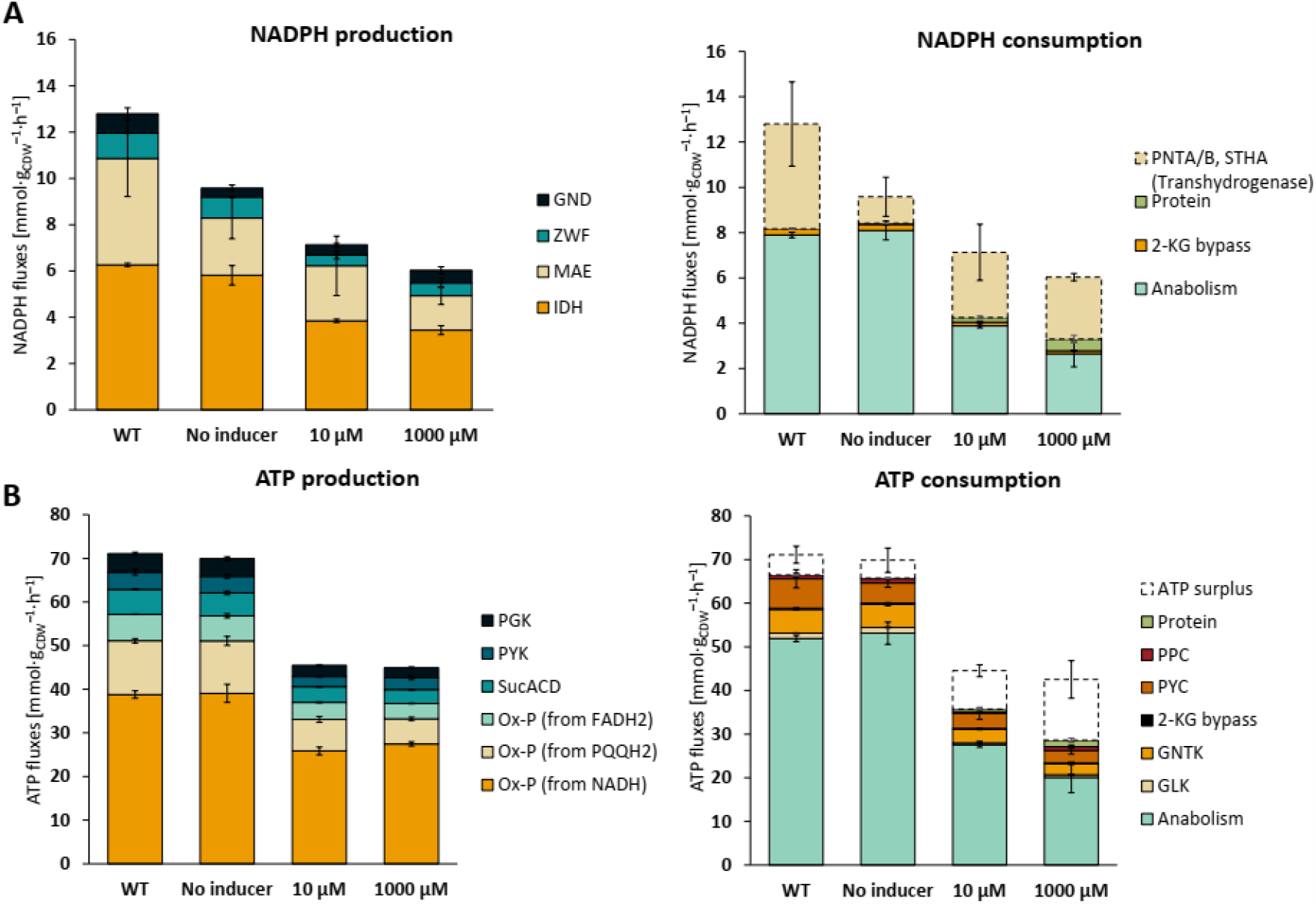
Redox and energy fluxes of *P. putida* KT2440 WT and *P. putida* CAP (pSEVA438_MBPeGFP) grown on glucose with or without 3-MB. The absolute fluxes (mmol·g_CDW_^−1^·h^−1^) related to the production and consumption of NADPH (A) and ATP (B) were determined from the carbon fluxes shown in Figure 4. Results are averages from 2 biological replicates for the WT strain and 3 biological replicates for the CAP (pSEVA438_MBPeGFP) strain; error bars represent standard deviations. Abbreviations: Gnd: 6-phosphogluconate dehydrogenase; Zwf: Glucose-6-P 1-dehydrogenase; Mae: malic enzyme; Idh: isocitrate deshydrogenase; Pgk: phosphoglycerate kinase; Pyk: pyruvate kinase; SucACD: succinyl CoA synthetase; Ox-P: oxidative phosphorylation; 2-KG bypass: 2-ketogluconate bypass; Ppc: phosphoenolpyruvate carboxylase; Pyc: pyruvate carboxylase; Gntk: gluconate kinase; Glk: Glucokinase.

Compared to the WT strain, NADPH production fluxes were respectively 27 %, 45 % and 54 % lower in the CAP (pSEVA438_MBPeGFP) strain under low, medium and high MBPeGFP production conditions, respectively (Figure 5A). While the NADPH consumption flux was similar to the WT strain’s under low MBPeGFP production conditions, it was much lower under medium and high MBPeGFP production. NADPH was mainly used for anabolism, and less than one-fifth (8.6%) of the total NADPH consumed was used for heterologous protein synthesis under high protein production conditions. As in the WT strain, NADPH production exceeded the cellular demand, with the excess amount accounting for a large proportion of total NADPH production under medium and high MBPeGFP production conditions (43 ± 11 % and 53 ± 5 % respectively). Excessive NADPH production during heterologous protein production has also been reported for other microorganisms (Daniels et al., 2018; Driouch et al., 2012; Nocon et al., 2016; Toya et al., 2014). Similar to the WT strain, the apparent surplus of NADPH was converted into NADH by transhydrogenase.

Regarding energy metabolism, low MBPeGFP production did not have an obvious impact on production and consumption fluxes in the CAP (pSEVA438_MBPeGFP) strain (Figure 5B). In contrast, under medium and high protein production conditions, ATP production and consumption fluxes were reduced by 36 % and 39 %, respectively, compared to the WT strain, while the contributions of oxidative phosphorylation and anabolic demand to the ATP balance remained close to the levels estimated in the WT strain. Remarkably, the amount of ATP used for heterologous protein production was very low, even under high induction. As observed for NADPH, a large apparent surplus of ATP was generated upon medium and high MBPeGFP production, representing respectively 25 % and 40 % of the total ATP consumed. This large apparent excess probably covered unquantified energy usages in protein biosynthesis, such as protein folding, as well as cellular adaptations to the stresses associated with protein overproduction (Hoffmann and Rinas, 2004).

### 3.7 Metabolic control analysis of heterologous protein production

We then aimed to quantify the extent of control exerted by heterologous protein production on metabolic fluxes using flux control coefficients 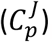, which quantify the degree of control exerted by a given parameter *p*, here the protein production rate, on each flux *J*. Control coefficients were calculated as the fractional change in the steady-state flux *J* in response to a fractional change in *p* (Fell and Thomas, 1995; Heinrich and Rapoport, 1974; Kacser and Burns, 1973):

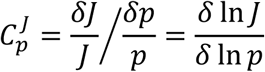

If the 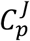 coefficients remain constant across different experiments in which *p* is modulated, their value can be estimated by fitting a linear function to a *ln*-*ln* plot, whose slope is equal to the control strength. Here, we used (steady-state) data collected for the various levels of protein production (Figure 4 and Table S2) to estimate the control exerted by protein production on carbon and energy fluxes. The 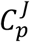 coefficients measure the sensitivity of the system to both direct metabolic regulation (though thermodynamics and metabolite-enzyme interactions), as well as to the indirect (hierarchical) action of signal transduction and gene expression.

The degree of control exerted by the MBPeGFP production rate on carbon and energy fluxes was investigated at the level of individual reactions and for more global processes such as growth and ATP and NADPH formation. This analysis provided an overall picture of the cellular processes affected by heterologous protein production (Figure 6). Among the analyzed fluxes, a majority of fluxes (25 out of 40, accounting for 63 %) showed a significant correlation with protein production (r^2^ > 0.5, p < 0.02). This indicates that the control exerted on these reactions remained stable across all MBPeGFP production conditions. The low r^2^ values (high p-values) obtained for the other fluxes may indicate an absence of control (*e*.*g*. Gnd) or reflect more complex, nonlinear control patterns that cannot be captured by linear regression (*e*.*g*. Ppc) (Figure S5).

**Fig. 6.**
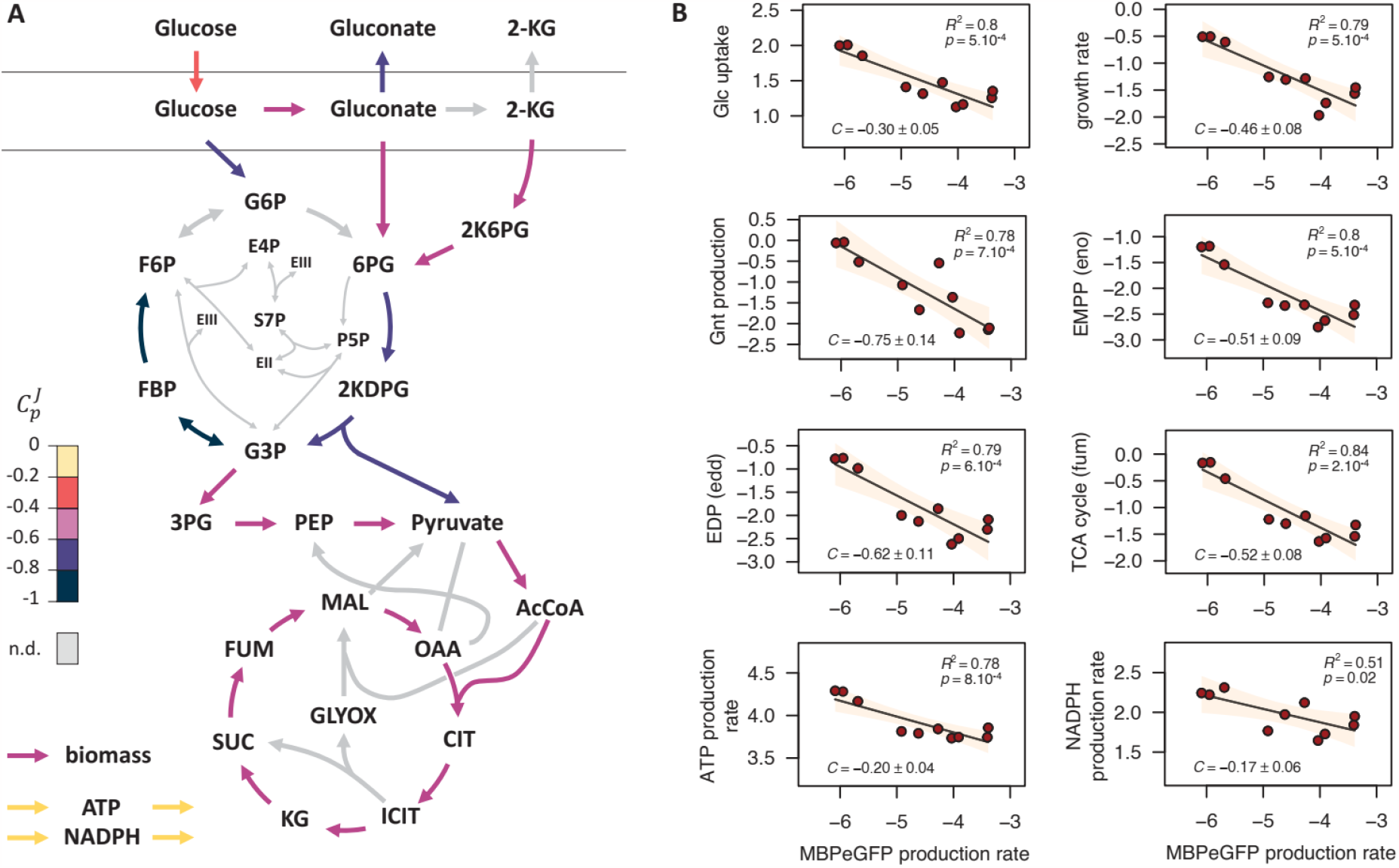
Control exerted on *P. putida*’s carbon and energy metabolism by heterologous protein production. Overview of the flux control coefficients (A). Control coefficients determined for carbon and energy fluxes through the main pathways (glucose uptake, EMPP, EDP, TCA cycle, ATP and NADPH production, and growth) (B), where dots represent experimental flux values (expressed on a logarithmic scale), lines represent the best linear fits, and shaded areas correspond to 95 % confidence intervals on the fits. Coefficients that could not be determined reliably for some reactions are marked *n*.*d*.

Looking more closely at the control coefficients, MBPeGFP production exerted a negative control on virtually all fluxes (with coefficients ranging from -0.17 to -0.99), in line with the global decrease in fluxes observed in response to MBPeGFP production. This negative control was observed for glucose uptake (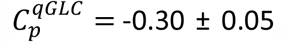 r^2^ = 0.80, p = 5.10^-4^), periplasmic glucose oxidation (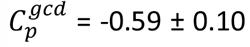, r^2^ = 0.80, p = 5.10^-4^), the ED pathway (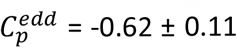, r^2^ = 0.79, p = 6.10^-4^), the EMP pathway (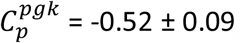, r^2^ = 0.80, p = 5.10^-4^; 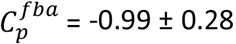, r^2^ = 0.60, p = 8.10^-3^), and the TCA cycle (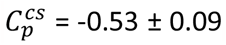, r^2^ = 0.83, p = 3.10^-4^). Significant control was also observed on ATP fluxes (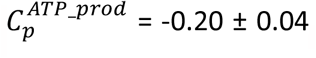, r^2^ = 0.78, p = 8.10^-4^), with similar levels of control on ATP production *via* oxidative phosphorylation and *via* substrate phosphorylation, and on NADPH fluxes (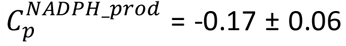 r^2^ = 0.51, p = 0.02). The control on energy and redox fluxes was weaker (coefficients between -0.17 and -0.19) than on carbon fluxes (coefficients between -0.30 and -0.99), indicating that in *P. putida* CAP (pSEVA438_MBPeGFP), energy fluxes are less sensitive to heterologous protein production than carbon fluxes are. These results also indicate that the fraction of glucose used for energy production increased with the level of heterologous protein production. These metabolic rearrangements were reflected in terms of cell physiology by a strong control on the growth rate (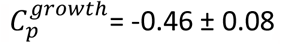; r^2^ = 0.79, p = 5.10^-4^).

These results offer a quantitative understanding of the impact of heterologous protein production on *P. putida*’s metabolism. The level of control remained largely stable across the three induction levels studied, indicating that all pathways responded smoothly to heterologous protein production. The fact that the response of carbon fluxes was more pronounced compared to energy fluxes highlights the flexibility of the central carbon metabolism and the robustness of the energy metabolism in *P. putida*.

## Conclusions

In this study, we conducted a detailed quantitative analysis to examine how *P. putida* responds to different degrees of metabolic burden caused by heterologous protein production. Our results indicate that low levels of heterologous protein production do not require any metabolic adaptation, because the extra demand does not exceed *P. putida*’s free capacity. However, when this free capacity is exceeded, growth is progressively inhibited and a global slowdown of metabolism is observed. Despite the relatively low (< 5%) carbon usage associated with heterologous protein production, our study revealed significant rearrangements in metabolic fluxes. These rearrangements indicate that even a small fraction of carbon dedicated to protein production can have a profound impact on the overall metabolic network of *P. putida*. As revealed by metabolic control analysis, heterologous protein production exerts a tighter control on carbon fluxes than on energy fluxes, suggesting that the flexible nature of *P. putida*’s central metabolic network is solicited to maintain energy production (Martin-Pascual et al., 2021; Tokic et al., 2020; Zobel et al., 2016). The metabolic flux response indicates a smooth, progressive decoupling of anabolism from catabolism, with energy production increasing far beyond the expected energy demands for protein biosynthesis. Similar behaviour was observed in other micro-organisms (Daniels et al., 2018; Driouch et al., 2012; Nocon et al., 2016; Toya et al., 2014), highlighting the genericity of microbial response to metabolic burdens. The reconfiguration of metabolic fluxes leads to an energy surplus well beyond what is necessary for growth and the production of heterologous proteins. This apparent surplus encompasses energy needs for various processes difficult to quantify and often underestimated. These include cellular maintenance, replication and expression of plasmids, the translation and folding of the heterologous protein, as well as the energy required to cope with the various other stresses induced by heterologous protein production. These stresses remain to be elucidated and warrant further investigation.

From a metabolic engineering perspective, this study showed that the expression of a very simple genetic circuit consisting of an inducible protomer and one gene encoding the MBP fused to eGFP, is accompanied by a drastic reduction in growth rate, which goes hand in hand with reduced glucose uptake, and a more pronounced conversion of glucose to gluconate and 2-KG, which then leaves the cell. The rearrangement of the metabolic fluxes is also reflected by less NADPH and ATP production. These findings could be a first starting point to streamline *P. putida* towards a better production of heterologous proteins following the design-build-test-learn (DBTL) cycle concept for metabolic engineering (Liu et al., 2015). In the design phase genetic targets will be selected, in the build phase the manipulation of *P. putida* will take place, which will then be characterized to obtain the data necessary to learn and predict parameters that can be applied in the next DBTL round. Interesting candidates that could be tackled in the first design phase is i) the improvement of glucose uptake by the introduction of an additional transporter, *e. g*. the Glf transporter (Bujdoš et al., 2023), ii) the blockage of the periplasmic glucose conversion to gluconate and 2-KG by deletion of *gcd* (Poblete-Castro et al., 2014) and, iii) the enhancement of the EDEMP cycle in order to modulate NADPH production by targeting the activity of *zwf* (Nikel et al., 2016a).

## Supporting information

Table S1 Network reaction model

Table S2 Linear stat

## 7. Acknowledgments

The authors thank MetaToul (Metabolomics & Fluxomics Facilities, Toulouse, France, www.metatoul.fr) and its staff for technical support and access to the NMR facility. MetaToul is part of the French National Infrastructure for Metabolomics and Fluxomics (www.metabohub.fr), funded by the ANR (MetaboHUB-ANR-11-INBS-0010). Edern Cahoreau is gratefully acknowledged for assistance with NMR spectroscopy. The French National Research Agency (ANR) and the German Research Foundation (DFG) are acknowledged for funding of the CHEAP project and the ANR for funding a post-doctoral fellowship to P.V.

