## Supplementary data for "Metabolic impact of heterologous protein production in *Pseudomonas putida*: Insights into carbon and energy flux control"

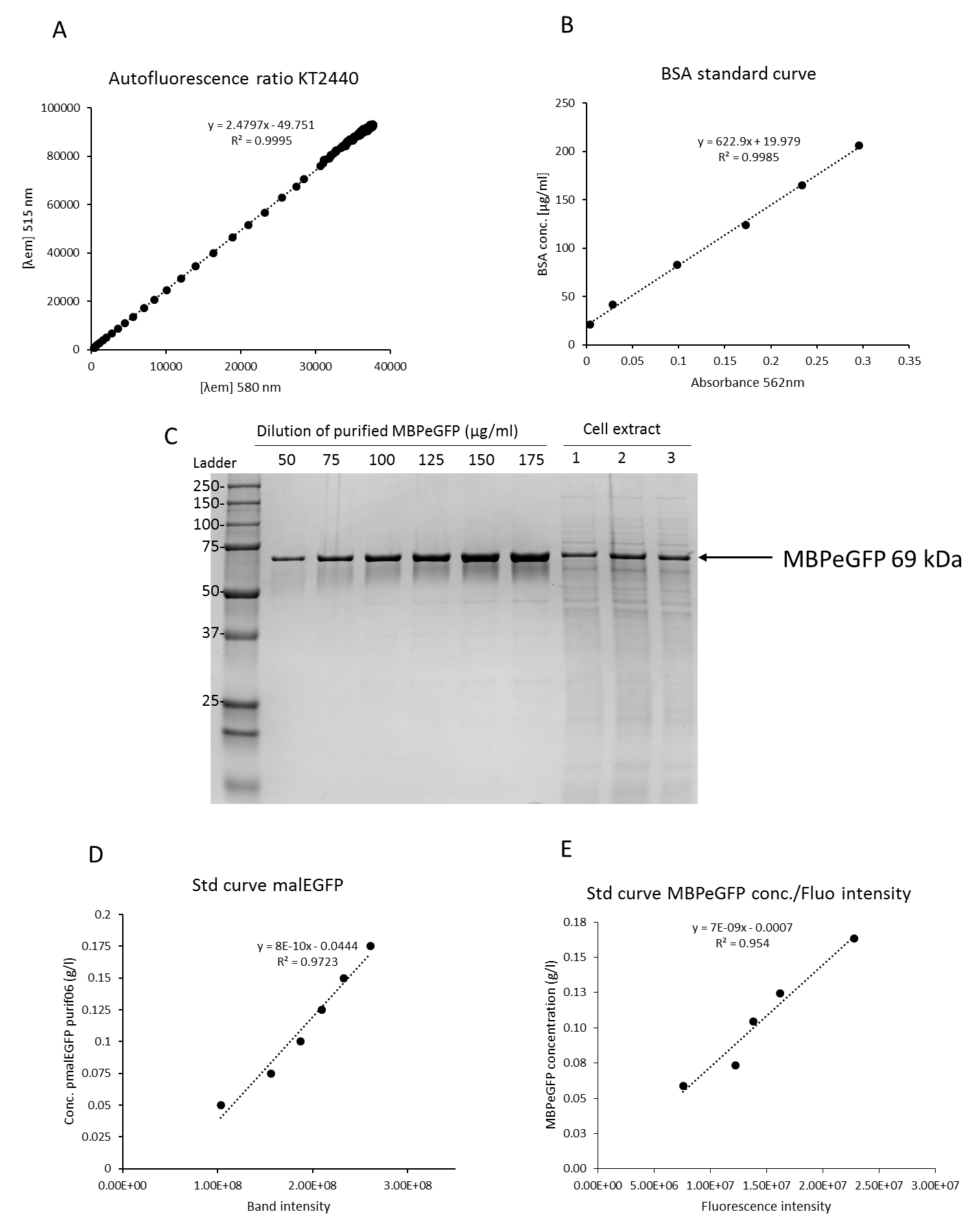

**Figure S1: Quantification of MBPeGFP concentration.** Standard curve used to determine the autofluorescence ratio. The ratio of autofluorescence *r*_a_ at 515 nm to that at 580 nm was calculated for a KT2440 WT strain culture grown in microtiter plate as described in the Materials & Methods section (A). BSA standard curve was construct by plotting BSA known concentrations against the absorbance read at 562 nm of BCA sample and slope of the curve was used to estimate the concentration of the purified MBPeGFP (B). SDS-PAGE containing different concentration of the MBPeGFP protein (175, 150, 125, 100, 75, 50 µg/mL) and diluted cellular extract of *P. putida* CAP-PROT cultivated with 1000 µM of inducer corresponding to 0.4 g_CDW_.l^-1^, lane 2: 0.5 g_CDW_.l^-1^ and lane 3: 0.35 g_CDW_.l^-1^ of cells (C). MBPeGFP standard curve obtained by plotting the MBPeGFP determined concentration against the band intensity. This standard curve was used to estimate the MBPeGFP concentration found in cellular extract (D). Green fluorescence intensity of the sample and the protein concentration estimated by SDS-PAGE at different time point of the culture were plotted to calculate the fluorescence factor (E).

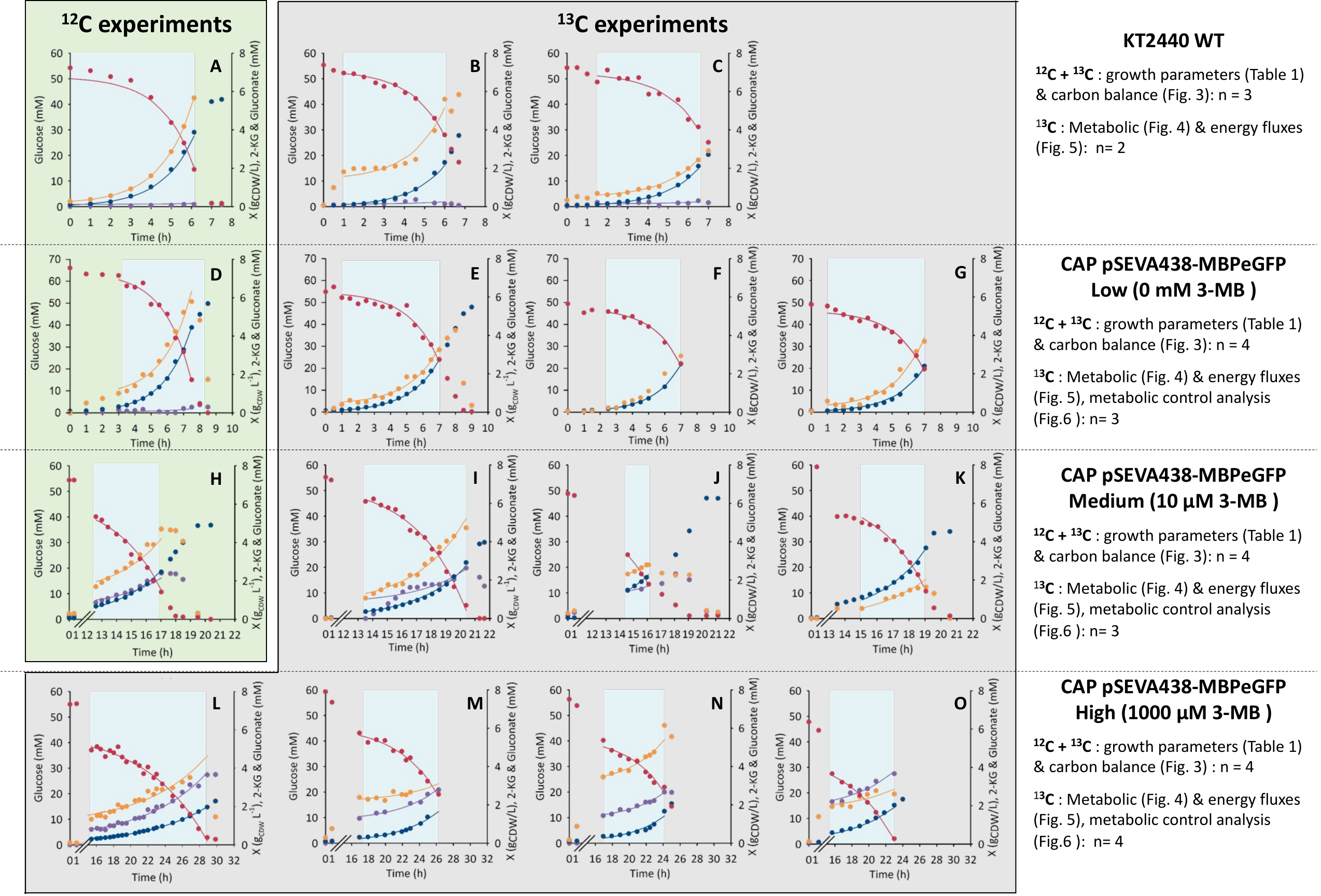

**Figure S2. Growth profiles of glucose-grown *P. putida* KT2440 wild-type (WT) strain (A, B, C) and *P. putida* CAP strain carrying pSEVA438_MBPeGFP during low (D, E, F, G), medium (H, I, J, K) and high (L, M, N, O) heterologous protein production**. Curves represent concentrations of cells (dark blue dots), glucose (red dots), gluconate (yellow dots) and ketogluconate (purple dots). The phase, during which samples were collected and growth parameters were calculated, is highlighted in blue. The solid lines represent the best fit in physiofit (Peiro *et al.*, 2019). The growth profiles in the green box derive from ^12^C glucose experiments and those in the grey box derive from ^13^C glucose experiments. The data from ^12^C and ^13^C experiments were used to determined growth parameters (Table 1) and carbon balances (Fig. 3). The data from ^13^C experiments were used to calculate metabolic fluxes (Fig. 4), redox & energy fluxes (Fig. 5) and to analyze metabolic control exerted by heterologous protein production on *P. putida*’s metabolism (Fig. 6). The right-hand column summarizes data use and indicates the number of biological replicates (n) used to construct Figures 3, 4, 5 and 6 and Table 1.

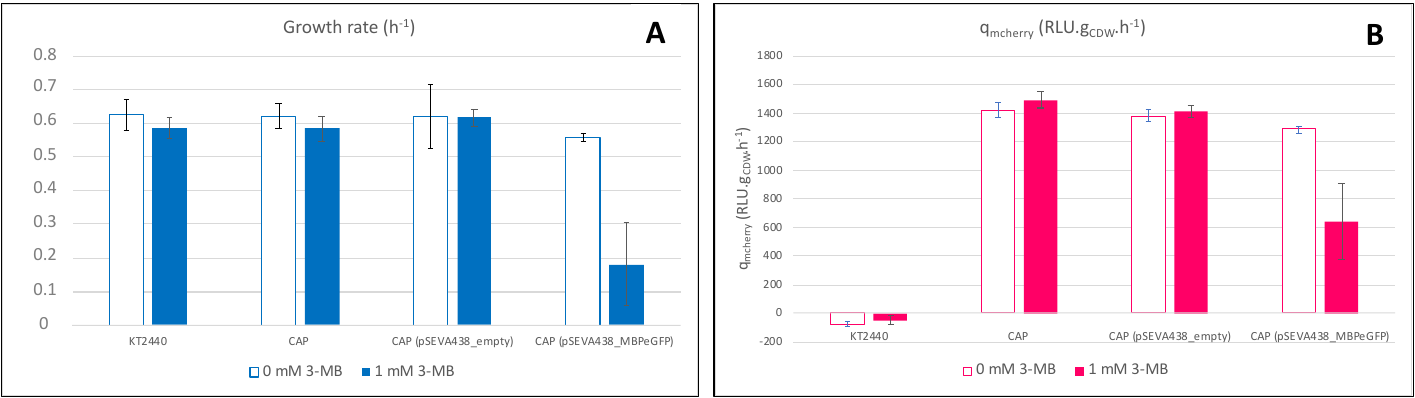

**Figure S3: Growth rate (A) and mCherry production rate** **(B) of *P. putida* KT2440 strain.** *P. putida* KT2440 wild-type strain, CAP strain, CAP strain carrying pSEVA438_empty or pSEVA438_MBPeGFP cultivated in microtiter plate in the presence and absence of 1 mM inducer (3-methyl-benzoate, 3-MB), performed in the same conditions as described in *Materials and Methods* section. Results are averages of two biological replicates.

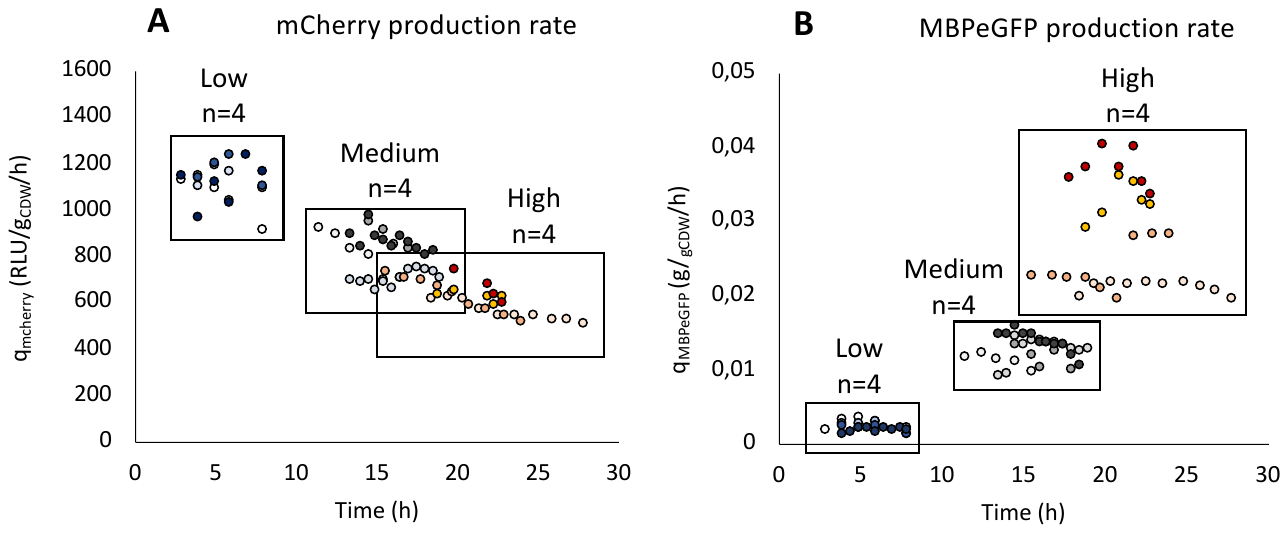

**Figure S4. mCherry (q_mcherry_) (A) and MBPeGFP (q_MBPeGFP_) (B) production rates during the exponential growth phase of glucose-grown *P. putida* CAP carrying pSEVA438_MBPEeGFP during low (blue), medium (grey) and high (orange) heterologous protein production.** Each biological replicate (n=4) is represented by different shades of color.

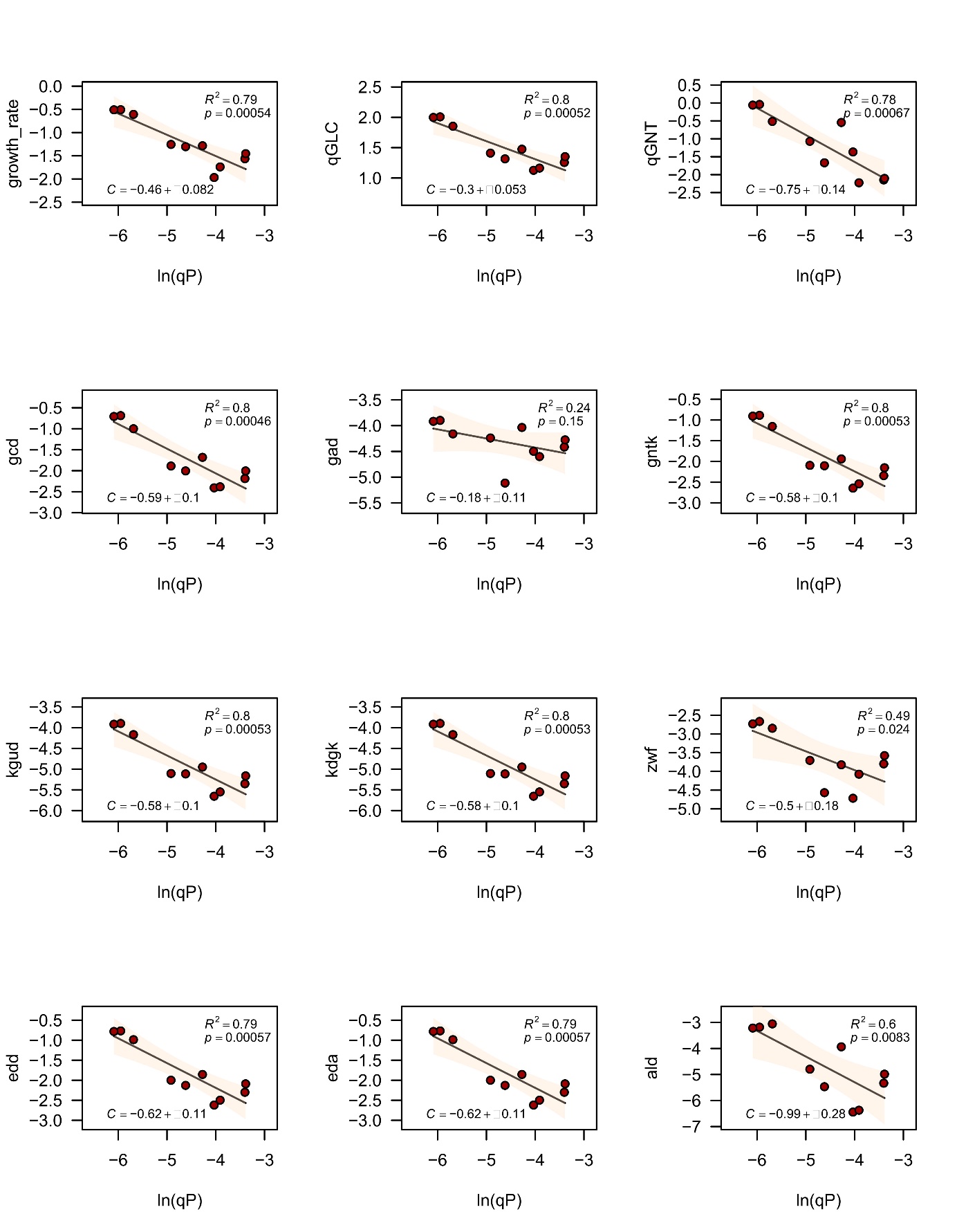

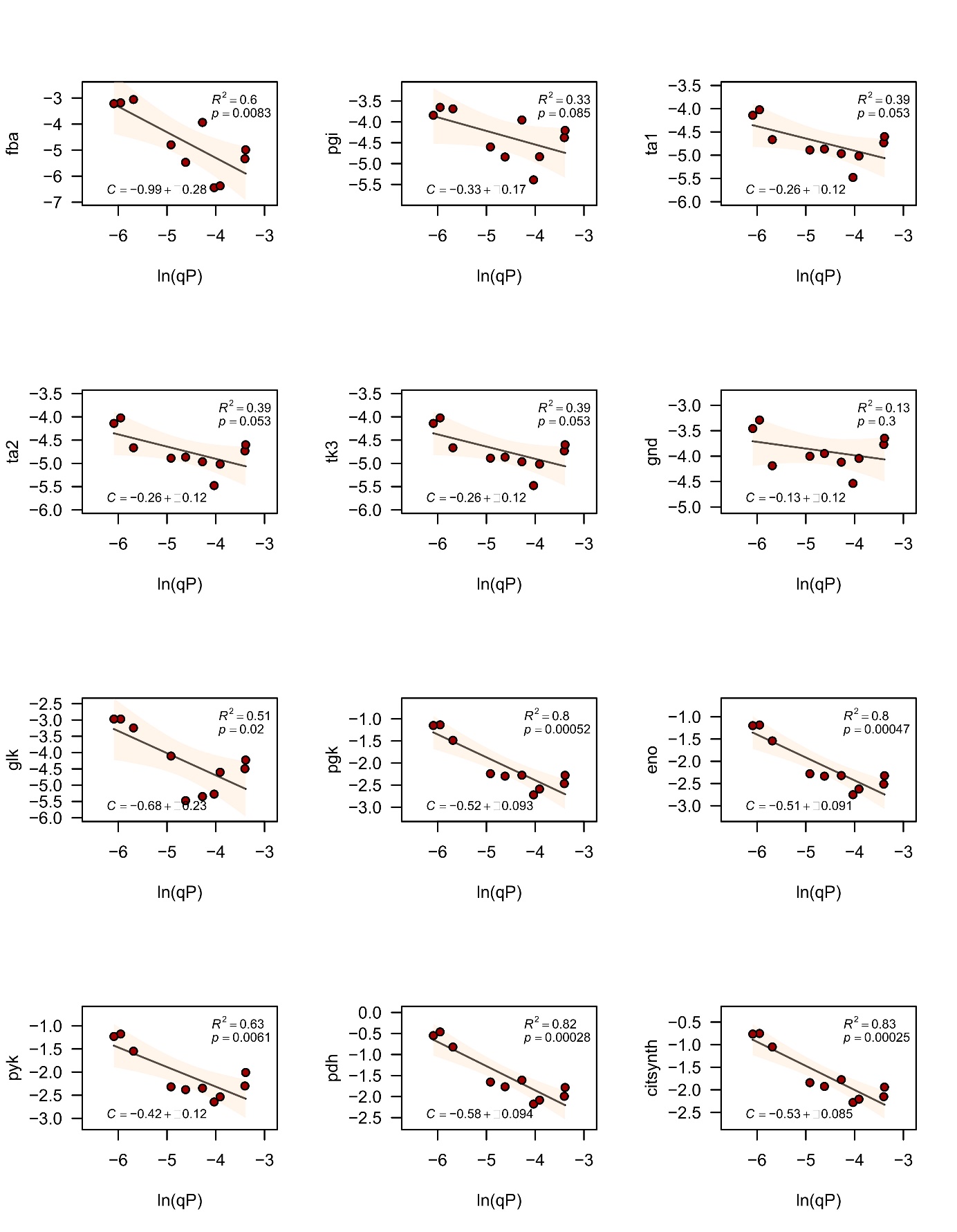

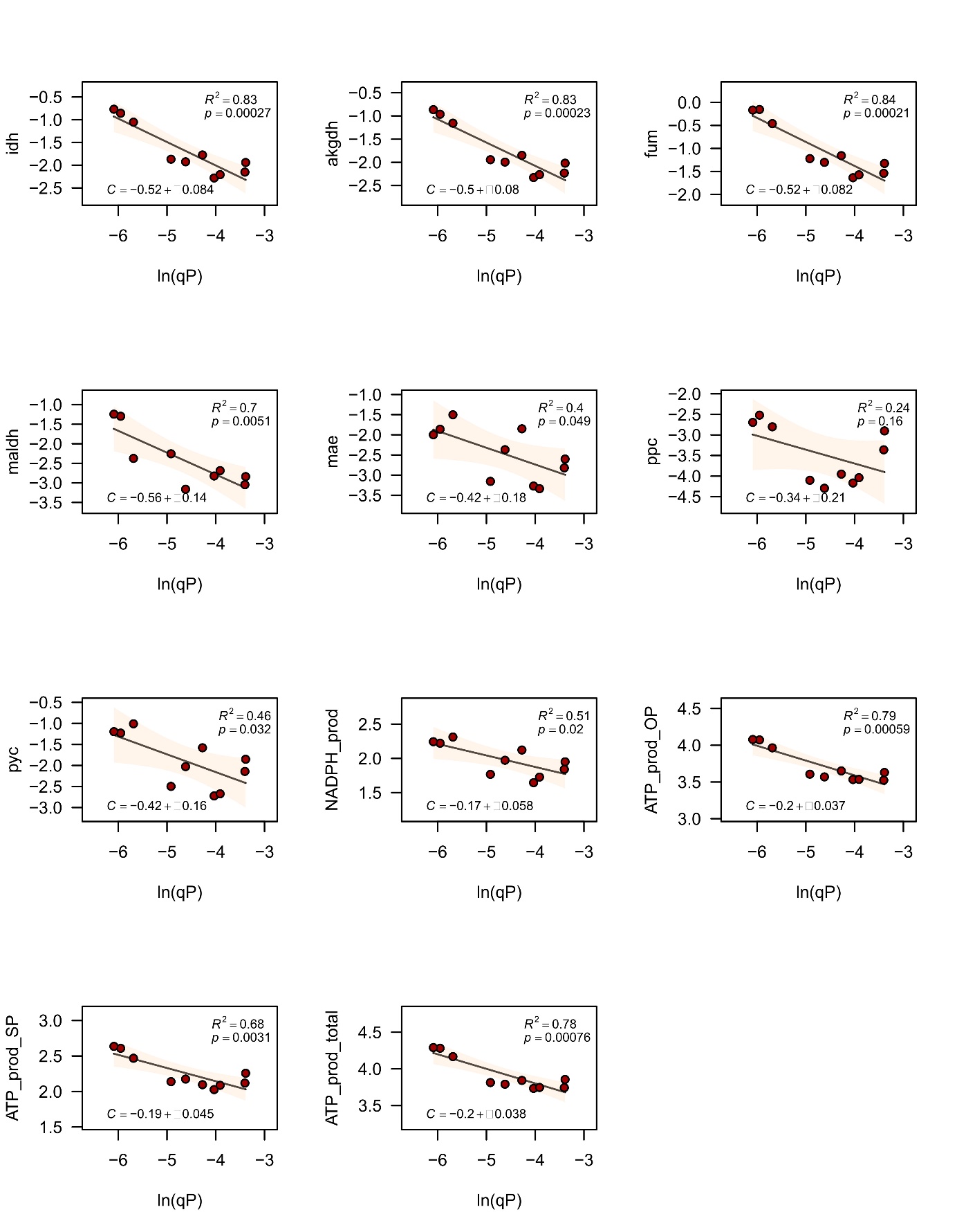

**Figure S5.** **Metabolic control analysis.** Control coefficients determined for all carbon and energy fluxes through the central metabolic pathways (glucose uptake, EMPP, EDP, TCA cycle, ATP and NADPH production, and growth). Dots represent experimental flux values (expressed in *ln* scale), lines represent the best linear fits, and shaded areas correspond to 95 % confidence intervals on the fit.

**A**

ATGAAAATCGAAGAAGGTAAACTGGTAATCTGGATTAACGGCGATAAAGGCTATAACGGTCTCGCTGAAGTCGGTAAGAAATTCGAGAAAGATACCGGAATTAAAGTCACCGTTGAGCATCCGGATAAACTGGAAGAGAAATTCCCACAGGTTGCGGCAACTGGCGATGGCCCTGACATTATCTTCTGGGCACACGACCGCTTTGGTGGCTACGCTCAATCTGGCCTGTTGGCTGAAATCACCCCGGACAAAGCGTTCCAGGACAAGCTGTATCCGTTTACCTGGGATGCCGTACGTTACAACGGCAAGCTGATTGCTTACCCGATCGCTGTTGAAGCGTTATCGCTGATTTATAACAAAGATCTGCTGCCGAACCCGCCAAAAACCTGGGAAGAGATCCCGGCGCTGGATAAAGAACTGAAAGCGAAAGGTAAGAGCGCGCTGATGTTCAACCTGCAAGAACCGTACTTCACCTGGCCGCTGATTGCTGCTGACGGGGGTTATGCGTTCAAGTATGAAAACGGCAAGTACGACATTAAAGACGTGGGCGTGGATAACGCTGGCGCGAAAGCGGGTCTGACCTTCCTGGTTGACCTGATTAAAAACAAACACATGAATGCAGACACCGATTACTCCATCGCAGAAGCTGCCTTTAATAAAGGCGAAACAGCGATGACCATCAACGGCCCGTGGGCATGGTCCAACATCGACACCAGCAAAGTGAATTATGGTGTAACGGTACTGCCGACCTTCAAGGGTCAACCATCCAAACCGTTCGTTGGCGTGCTGAGCGCAGGTATTAACGCCGCCAGTCCGAACAAAGAGCTGGCGAAAGAGTTCCTCGAAAACTATCTGCTGACTGATGAAGGTCTGGAAGCGGTTAATAAAGACAAACCGCTGGGTGCCGTAGCGCTGAAGTCTTACGAGGAAGAGTTGGCGAAAGATCCACGTATTGCCGCCACCATGGAAAACGCCCAGAAAGGTGAAATCATGCCGAACATCCCGCAGATGTCCGCTTTCTGGTATGCCGTGCGTACTGCGGTGATCAACGCCGCCAGCGGTCGTCAGACTGTCGATGAAGCCCTGAAAGACGCGCAGACTCGTATCACCAAGggtggtggtggttcgATGGTGAGCAAGGGCGAGGAGCTGTTCACCGGGGTGGTGCCCATCCTGGTCGAGCTGGACGGCGACGTAAACGGCCACAAGTTCAGCGTGTCCGGCGAGGGCGAGGGCGATGCCACCTACGGCAAGCTGACCCTGAAGTTCATCTGCACCACCGGCAAGCTGCCCGTGCCCTGGCCCACCCTCGTGACCACCCTGACCTACGGCGTGCAGTGCTTCAGCCGCTACCCCGACCACATGAAGCAGCACGACTTCTTCAAGTCCGCCATGCCCGAAGGCTACGTCCAGGAGCGCACCATCTTCTTCAAGGACGACGGCAACTACAAGACCCGCGCCGAGGTGAAGTTCGAGGGCGACACCCTGGTGAACCGCATCGAGCTGAAGGGCATCGACTTCAAGGAGGACGGCAACATCCTGGGGCACAAGCTGGAGTACAACTACAACAGCCACAACGTCTATATCATGGCCGACAAGCAGAAGAACGGCATCAAGGTGAACTTCAAGATCCGCCACAACATCGAGGACGGCAGCGTGCAGCTCGCCGACCACTACCAGCAGAACACCCCCATCGGCGACGGCCCCGTGCTGCTGCCCGACAACCACTACCTGAGCACCCAGTCCGCCCTGAGCAAAGACCCCAACGAGAAGCGCGATCACATGGTCCTGCTGGAGTTCGTGACCGCCGCCGGGATCACTCTCGGCATGGACGAGCTGTACAAGGCTAGCCATCATCATCATCATCACTAA

**B**

MKIEEGKLVIWINGDKGYNGLAEVGKKFEKDTGIKVTVEHPDKLEEKFPQVAATGDGPDIIFWAHDRFGGYAQSGLLAEITPDKAFQDKLYPFTWDAVRYNGKLIAYPIAVEALSLIYNKDLLPNPPKTWEEIPALDKELKAKGKSALMFNLQEPYFTWPLIAADGGYAFKYENGKYDIKDVGVDNAGAKAGLTFLVDLIKNKHMNADTDYSIAEAAFNKGETAMTINGPWAWSNIDTSKVNYGVTVLPTFKGQPSKPFVGVLSAGINAASPNKELAKEFLENYLLTDEGLEAVNKDKPLGAVALKSYEEELAKDPRIAATMENAQKGEIMPNIPQMSAFWYAVRTAVINAASGRQTVDEALKDAQTRITKggggsMVSKGEELFTGVVPILVELDGDVNGHKFSVSGEGEGDATYGKLTLKFICTTGKLPVPWPTLVTTLTYGVQCFSRYPDHMKQHDFFKSAMPEGYVQERTIFFKDDGNYKTRAEVKFEGDTLVNRIELKGIDFKEDGNILGHKLEYNYNSHNVYIMADKQKNGIKVNFKIRHNIEDGSVQLADHYQQNTPIGDGPVLLPDNHYLSTQSALSKDPNEKRDHMVLLEFVTAAGITLGMDELYKASHHHHHH

**Nucleotide (A) and amino acid (B) sequences of MBPeGFP.** *malE* sequences is underlined in gray, *eGFP* in green and the linker in cyan. his-tag is indicated in red font.
