## Supplementary material for "Metabolic impact of heterologous protein production in *Pseudomonas putida*: Insights into carbon and energy flux control": Table S1 Network reaction model

| **Pathway** | **Gene / function** | **Model reaction name** | **Reaction equation (= reversible, > irreversible)** |
| --- | --- | --- | --- |
| Periplasmic pathway / Uptake substrate | *oprB* | Glc_in | GLC_EX > GLC |
|  | *gcd* | Gcd | GLC > GNT + PQQH2 |
|  | Gluconate secretion | Out_Gnt | GNT > GNT_EX |
|  | *gntP, gnuK* | Gntk | GNT + ATP > 6PGNT |
|  | *gad* | Gad | GNT > 2KGNT + FADH2 |
|  | 2K-gluconate secretion | Out_2-KG | 2KGNT > 2KGNT_EX |
|  | *kguT, kguK* | Kdgk | 2KGNT + ATP > 2K6PG |
|  | *kguD* | Kgud | 2K6PG + NADPH > 6PGNT |
| Embden Meyerhof Parnas pathway | *gtsABCD, glk* | Glk | GLC + 2 ATP > G6P |
|  | *pgi-1, 2* | Pgi | F6P = G6P |
|  | *fba* | Fbp | FBP > F6P + Pi |
|  | *fda, PP_1791, PP_2871, PP_3224* | Ald | DHAP + GA3P = FBP |
|  | *gapA, gapB, epd, PP - 0665, PP - 3443 pgk* | Pgk | GA3P = 3PG + NADH + ATP |
|  | *pgm, PP_2243, PP_3923, PP_4450, eno* | Eno | 3PG = PEP |
|  | *pykA, pykF* | Pyk | PEP > PYR + ATP |
| Entner-Doudoroff pathway | *edd* | Edd | 6PGNT > 2KDPG |
|  | *eda* | Eda | 2KDPG > PYR + GA3P |
| Pentose phosphate pathway | *zwf, zwfA, zwfB* | Zwf | G6P > 6PGNT + NADPH |
|  | *gnd, rpe* | Gnd | 6PGNT > P5P + CO2 + NADPH |
|  | *tktA* | Tk1 | P5P = GA3P + E2 |
|  | *tktA* | Tk2 | E4P + E2 = F6P |
|  | *tktA* | Tk3 | P5P = GA3P + E4P |
|  | *tal* | Ta1 | GA3P + E3 = S7P |
|  | *tal* | Ta2 | S7P + GA3P = E4P + F6P |
| TCA cycle / Glyoxylate shunt | *acoABC, aceEF, IpdGV, Ipd3* | Pdh | PYR > ACCOA + CO2 + NADH |
|  | *gltA, acnAI-II acnB* | Citsynth | ACCOA + OAA > ICIT |
|  | *icd, idh* | Idh | ICIT > AKG + CO2 + NADPH |
|  | *Lpd, LpdGV, sucABCD, scpC, PP_2652, PP_3662* | aKgdh/SucACD | AKG = SUC + CO2 + ATP + NADH |
|  | *sdhABCD, kgdB* | Fum_a | SUC = FUM + FADH2 |
|  | *fumC - 1, 2, PP_0897* | Fum_b | FUM = MAL |
|  | *mdh, PP_3591, mqo-1, 2, 3* | MaldH | MAL = OAA + NADH |
|  | *aceA* | AceA | ICIT > SUC + GLYOX |
|  | *glcB* | AceB | ACCOA + GLYOX > MAL |
| Anaplerotic pathway | *maeB* | Mae | MAL > PYR + CO2 + NADPH |
|  | *accC-2, oadA* | Pyc | PYR + CO2 + ATP > OAA |
|  | *ppc* | Ppc | OAA + ATP > PEP + CO2 |
| Biomass synthesis | growth | bs_glc6P | G6P > |
|  |  | bs_fru6P | F6P > |
|  |  | bs_ga3p | GA3P > |
|  |  | bs_pga | 3PG > |
|  |  | bs_pyr | PYR > |
|  |  | bs_e4p | E4P > |
|  |  | bs_rib5p | P5P > |
|  |  | bs_pep | PEP > |
|  |  | bs_accoa | ACCOA > |
|  |  | bs_akg | AKG > |
|  |  | bs_oaa | OAA > |
| Protein synthesis | protein production | bsp_rib5p | P5P > |
|  |  | bsp_e4p | E4P > |
|  |  | bsp_pga | 3PG > |
|  |  | bsp_pep | PEP > |
|  |  | bsp_pyr | PYR > |
|  |  | bsp_accoa | ACCOA > |
|  |  | bsp_oaa | OAA > |
|  |  | bsp_akg | AKG > |
