## Supplementary material for "Metabolic impact of heterologous protein production in *Pseudomonas putida*: Insights into carbon and energy flux control": Table S2 Linear stat

**Table S2 Rates (A) and linear stat (B)**

**A Rates**

|  | **growth rate (h-^1^)** | **qglucose (mmol · [g_CDW_ · h]^-1^)** | **qKG (mmol · [g_CDW_ · h]^-1^)** | **qGlc (mmol · [g_CDW_ · h]^-1^)** | **qprot (g_prot_ · [g_CDW_ · h]^-1^)** | **BM/qGlc (g_CDW_/mmol_Glc)** | **qKG/qGlc (mmol_KG/mmol_Glc)** | **qGlnt/qGlc (mmol_Glnt/mmol_Glc)** | **qprot/mmol_Glc (gProt/mmol**  **_Glc)** |
| --- | --- | --- | --- | --- | --- | --- | --- | --- | --- |
| **WT_rep01** | 0.5582 | 7.3474 | 0.0892 | 1.0132 | ND | 0.0760 | 0.0121 | 0.1379 | ND |
| **WT_rep02** | 0.5700 | 7.0229 | 0.0330 | 0.5695 | ND | 0.0812 | 0.0047 | 0.0811 | ND |
| **CAP (pSEVA438_MBPeGFP)_rep01** | 0.6020 | 7.4457 | 0.0000 | 0.9597 | 0.0026 | 0.0809 | 0.0000 | 0.1289 | 0.0004 |
| **CAP (pSEVA438_MBPeGFP)_rep02** | 0.6010 | 7.3771 | 0.0000 | 0.9416 | 0.0023 | 0.0815 | 0.0000 | 0.1276 | 0.0003 |
| **CAP (pSEVA438_MBPeGFP)_rep03** | 0.5445 | 6.3823 | 0.0000 | 0.5966 | 0.0034 | 0.0853 | 0.0000 | 0.0935 | 0.0005 |
| **10µM_CAP (pSEVA438_MBPeGFP)_rep01** | 0.2849 | 4.0979 | 0.2043 | 0.3428 | 0.0073 | 0.0695 | 0.0499 | 0.0837 | 0.0018 |
| **10µM_CAP (pSEVA438_MBPeGFP)_rep02** | 0.2769 | 4.3699 | 0.2426 | 0.5800 | 0.0140 | 0.0634 | 0.0555 | 0.1327 | 0.0032 |
| **10µM_CAP (pSEVA438_MBPeGFP)_rep03** | 0.2713 | 3.7289 | 0.0000 | 0.1885 | 0.0099 | 0.0728 | 0.0000 | 0.0505 | 0.0026 |
| **1mM_CAP (pSEVA438_MBPeGFP)_rep01** | 0.1397 | 3.0855 | 0.2474 | 0.2549 | 0.0177 | 0.0453 | 0.0802 | 0.0826 | 0.0057 |
| **1mM_CAP (pSEVA438_MBPeGFP)_rep02** | 0.1755 | 3.2003 | 0.1926 | 0.1079 | 0.0200 | 0.0548 | 0.0602 | 0.0337 | 0.0063 |
| **1mM_CAP (pSEVA438_MBPeGFP)_rep03** | 0.2092 | 3.5128 | 0.2092 | 0.1168 | 0.0333 | 0.0595 | 0.0595 | 0.0333 | 0.0095 |
| **1mM_CAP (pSEVA438_MBPeGFP)_rep04** | 0.2334 | 3.8650 | 0.2110 | 0.1219 | 0.0338 | 0.0604 | 0.0546 | 0.0315 | 0.0087 |

**B Linear stat**

|  | WT_rep01 | WT_rep02 | CAP (pSEVA438_MBPeGFP)_rep01 | CAP (pSEVA438_MBPeGFP)_rep02 | CAP (pSEVA438_MBPeGFP)_rep03 | 10µM_CAP (pSEVA438_MBPeGFP)_rep01 | 10µM_CAP (pSEVA438_MBPeGFP)_rep02 | 10µM_CAP (pSEVA438_MBPeGFP)_rep03 | 1mM_CAP (pSEVA438_MBPeGFP)_rep01 | 1mM_CAP (pSEVA438_MBPeGFP)_rep02 | 1mM_CAP (pSEVA438_MBPeGFP)_rep03 | 1mM_CAP (pSEVA438_MBPeGFP)_rep04 |
| --- | --- | --- | --- | --- | --- | --- | --- | --- | --- | --- | --- | --- |
| row_col | value | value | value | value | value | value | value | value | value | value | value | value |
| d.n.aceA | 1.00E-04 | 1.00E-04 | 0.085291777 | 0.004536037 | 0.002893288 | 0.025677824 | 1.00E-04 | 1.00E-04 | 0.0001 | 1.00E-04 | 1.00E-04 | 1.00E-04 |
| d.n.aceB | 0.0001 | 1.00E-04 | 0.085291777 | 0.004536037 | 0.002893288 | 0.025677824 | 0.0001 | 1.00E-04 | 0.0001 | 0.0001 | 0.0001 | 0.0001 |
| d.n.akgdh | 0.772621992 | 0.821094723 | 0.6879767 | 0.773715324 | 0.772420188 | 0.852213462 | 0.823555437 | 0.977563354 | 1.022944246 | 1.011975208 | 0.868626357 | 0.888073036 |
| d.n.ald | 0.088878291 | 0.056252531 | 0.074842577 | 0.073708284 | 0.11550973 | 0.049183964 | 0.102246586 | 0.030316698 | 0.016734766 | 0.016673682 | 0.039131826 | 0.045675599 |
| d.n.bs_accoa | 0.191139848 | 0.204228386 | 0.200672567 | 0.20095137 | 0.217437019 | 0.168209107 | 0.159080441 | 0.178628927 | 0.113057843 | 0.136793442 | 0.15515726 | 0.15733816 |
| d.n.bs_akg | 0.073041496 | 0.078043103 | 0.076684295 | 0.076790836 | 0.083090603 | 0.064278825 | 0.060790429 | 0.068260618 | 0.043203519 | 0.052273755 | 0.059291239 | 0.06012464 |
| d.n.bs_e4p | 0.082650134 | 0.088309705 | 0.086772145 | 0.086892702 | 0.094021205 | 0.07273473 | 0.068787435 | 0.07724033 | 0.048886959 | 0.059150389 | 0.067091025 | 0.068034061 |
| d.n.bs_fru6P | 0.005224202 | 0.005581936 | 0.005484749 | 0.005492369 | 0.005942952 | 0.004597463 | 0.00434796 | 0.004882256 | 0.003090078 | 0.003738815 | 0.004240732 | 0.00430034 |
| d.n.bs_ga3p | 0.05102651 | 0.054520614 | 0.053571355 | 0.053645785 | 0.058046777 | 0.044904941 | 0.042467961 | 0.047686607 | 0.030181813 | 0.036518246 | 0.041420633 | 0.042002844 |
| d.n.bs_glc6P | 0.013246113 | 0.014153158 | 0.013906737 | 0.013926058 | 0.015068524 | 0.011656998 | 0.011024376 | 0.012379099 | 0.007834981 | 0.009479873 | 0.010752497 | 0.010903635 |
| d.n.bs_oaa | 0.102986344 | 0.110038462 | 0.108122583 | 0.108272802 | 0.117155286 | 0.090631238 | 0.085712703 | 0.096245447 | 0.060915681 | 0.073704445 | 0.083598889 | 0.08477396 |
| d.n.bs_pep | 0.133863007 | 0.143029443 | 0.140539158 | 0.140734416 | 0.152279985 | 0.117803677 | 0.111410501 | 0.125101101 | 0.079179005 | 0.09580201 | 0.108662938 | 0.110190311 |
| d.n.bs_pga | 0.025839579 | 0.027608976 | 0.027128276 | 0.027165967 | 0.029394609 | 0.022739646 | 0.021505571 | 0.024148268 | 0.015283925 | 0.018492664 | 0.020975209 | 0.021270038 |
| d.n.bs_pyr | 0.008209029 | 0.008771153 | 0.008618438 | 0.008630412 | 0.009338434 | 0.007224205 | 0.006832149 | 0.007671713 | 0.004855582 | 0.005874972 | 0.006663657 | 0.006757322 |
| d.n.bs_rib5p | 0.054478376 | 0.058208851 | 0.057195377 | 0.057274841 | 0.061973554 | 0.047942693 | 0.045340855 | 0.050912534 | 0.032223567 | 0.03898865 | 0.044222676 | 0.044844273 |
| d.n.bsp_accoa | 7.33E-06 | 7.33E-06 | 0.000256553 | 0.000225762 | 0.000187675 | 0.001368896 | 0.001391869 | 0.001714045 | 0.003784261 | 0.004578207 | 0.006997727 | 0.006451244 |
| d.n.bsp_akg | 1.51E-05 | 1.51E-05 | 0.000528225 | 0.000464827 | 0.00038641 | 0.002818458 | 0.002865757 | 0.003529095 | 0.007791521 | 0.009426197 | 0.014407814 | 0.013282646 |
| d.n.bsp_e4p | 8.91E-06 | 8.91E-06 | 0.000311916 | 0.00027448 | 0.000228175 | 0.001664298 | 0.001692228 | 0.002083929 | 0.004600889 | 0.005566165 | 0.008507808 | 0.007843396 |
| d.n.bsp_oaa | 3.13E-05 | 3.13E-05 | 0.001096695 | 0.000965069 | 0.00080226 | 0.005851653 | 0.005949855 | 0.007327069 | 0.016176671 | 0.019570569 | 0.029913348 | 0.027577286 |
| d.n.bsp_pep | 1.65E-05 | 1.65E-05 | 0.000578549 | 0.000509111 | 0.000423223 | 0.003086972 | 0.003138777 | 0.003865311 | 0.008533816 | 0.010324227 | 0.015780441 | 0.014548079 |
| d.n.bsp_pga | 1.16E-05 | 1.16E-05 | 0.00040749 | 0.000358583 | 0.000298089 | 0.00217425 | 0.002210738 | 0.002722458 | 0.006010631 | 0.007271673 | 0.011114654 | 0.010246663 |
| d.n.bsp_pyr | 4.66E-05 | 4.66E-05 | 0.001629959 | 0.001434331 | 0.001192356 | 0.008696999 | 0.008842952 | 0.010889832 | 0.024042525 | 0.029086694 | 0.044458616 | 0.040986651 |
| d.n.bsp_rib5p | 4.03E-06 | 4.03E-06 | 0.000140858 | 0.000123952 | 0.000103041 | 0.000751576 | 0.000764189 | 0.000941076 | 0.002077704 | 0.002513611 | 0.00384202 | 0.00354198 |
| d.n.citsynth | 0.845778582 | 0.899252921 | 0.850480997 | 0.855507024 | 0.858790487 | 0.944988569 | 0.887311623 | 1.049453067 | 1.074039286 | 1.073775161 | 0.94242541 | 0.961580322 |
| d.n.eda | 0.826402318 | 0.833889797 | 0.837387931 | 0.840507611 | 0.91547926 | 0.806338747 | 0.818971911 | 0.855917673 | 0.765049375 | 0.803419815 | 0.811702615 | 0.828529827 |
| d.n.edd | 0.826402318 | 0.833889797 | 0.837387931 | 0.840507611 | 0.91547926 | 0.806338747 | 0.818971911 | 0.855917673 | 0.765049375 | 0.803419815 | 0.811702615 | 0.828529827 |
| d.n.eno | 0.52873286 | 0.609590542 | 0.551820088 | 0.553978349 | 0.525604477 | 0.608619851 | 0.514355146 | 0.696768552 | 0.670530335 | 0.707789864 | 0.655654724 | 0.655080793 |
| d.n.fba | 0.088878291 | 0.056252531 | 0.074842577 | 0.073708284 | 0.11550973 | 0.049183964 | 0.102246586 | 0.030316698 | 0.016734766 | 0.016673682 | 0.039131826 | 0.045675599 |
| d.n.fum_a | 0.386360996 | 0.410597362 | 0.386634238 | 0.38912568 | 0.387656738 | 0.438945643 | 0.411827718 | 0.488831677 | 0.511522123 | 0.506037604 | 0.434363179 | 0.444086518 |
| d.n.fum_b | 0.386360996 | 0.410597362 | 0.386634238 | 0.38912568 | 0.387656738 | 0.438945643 | 0.411827718 | 0.488831677 | 0.511522123 | 0.506037604 | 0.434363179 | 0.444086518 |
| d.n.gad | 0.04827204 | 0.043392164 | 0.036582528 | 0.036559476 | 0.038313242 | 0.085826211 | 0.092558361 | 0.043189382 | 0.116867241 | 0.098066434 | 0.098037973 | 0.093033484 |
| d.n.gcd | 0.91951491 | 0.909918175 | 0.907626273 | 0.905875467 | 0.904224709 | 0.901952606 | 0.975092291 | 0.969852037 | 0.946198984 | 0.902143177 | 0.909656523 | 0.902390516 |
| d.n.Glc_in_U | 0.177589255 | 0.17979355 | 0.174771707 | 0.17810649 | 0.175068856 | 0.167174006 | 0.169693799 | 0.168396685 | 0.176836316 | 0.180558507 | 0.177013285 | 0.177887401 |
| d.n.gntk | 0.733483371 | 0.785475685 | 0.742625314 | 0.74215736 | 0.772368149 | 0.732823346 | 0.752753866 | 0.87674446 | 0.746863336 | 0.770377194 | 0.778271815 | 0.777817981 |
| d.n.idh | 0.845678582 | 0.899152921 | 0.765189221 | 0.850970987 | 0.8558972 | 0.919310746 | 0.887211623 | 1.049353067 | 1.073939286 | 1.073675161 | 0.94232541 | 0.961480322 |
| d.n.kgud | 0.036132186 | 0.038693384 | 0.036582528 | 0.036559476 | 0.038047692 | 0.036099672 | 0.037081471 | 0.043189382 | 0.036791297 | 0.037949615 | 0.038338513 | 0.038316157 |
| d.n.mae | 0.46730974 | 0.821094723 | 0.280538899 | 0.24922069 | 0.546629433 | 0.253812259 | 0.823555437 | 0.673359823 | 0.399043551 | 0.347259307 | 0.483206677 | 0.496676851 |
| d.n.maldh | 0.305412252 | 1.00E-04 | 0.492729578 | 0.52903067 | 0.228684042 | 6.24E-01 | 1.00E-04 | 0.304303531 | 0.624000695 | 0.664815901 | 0.38551968 | 0.391496185 |
| d.n.out_co2 | 2.869685401 | 3.317571602 | 7.596203713 | 6.351980075 | 2.546312393 | 6.644872072 | 1000 | 4.919204665 | 3.892998435 | 5.16544122 | 2.91496201 | 2.963770844 |
| d.n.out_Gnt | 0.137759499 | 0.081050326 | 0.128418431 | 0.127158631 | 0.093543318 | 0.083303049 | 0.129780064 | 0.049918194 | 0.082468407 | 0.033699549 | 0.033346736 | 0.031539051 |
| d.n.pgi | 0.080242847 | 0.079683294 | 0.046890757 | 0.039498378 | 0.06158446 | 0.059921268 | 0.100499645 | 0.056841917 | 0.047986261 | 0.077651412 | 0.101942444 | 0.100096132 |
| d.n.pgk | 5.55E-01 | 6.37E-01 | 0.579355854 | 0.581502899 | 0.555297175 | 0.633533747 | 0.538071455 | 0.723639278 | 0.691824891 | 0.733554201 | 0.687744587 | 0.686597493 |
| d.n.ppc | 0.076651621 | 0.16388291 | 0.145133953 | 0.124096184 | 0.149076517 | 0.098517492 | 0.100605012 | 0.097996868 | 0.162709772 | 0.171970376 | 0.281200095 | 0.367652696 |
| d.n.ta1 | 0.039623903 | 0.058665659 | 0.032308495 | 0.029224822 | 0.023133531 | 0.044866898 | 0.036540341 | 0.055365867 | 0.04391471 | 0.06471655 | 0.071325091 | 0.067299165 |
| d.n.ta2 | 0.039623903 | 0.058665659 | 0.032308495 | 0.029224822 | 0.023133531 | 0.044866898 | 0.036540341 | 0.055365867 | 0.04391471 | 0.06471655 | 0.071325091 | 0.067299165 |
| d.n.tk1 | -0.003411241 | 0.029012699 | -0.022467072 | -0.028717537 | -0.047982318 | 0.015334767 | 2.60E-03 | 0.031407474 | 0.034341572 | 0.064716546 | 0.067051349 | 0.058720873 |
| d.n.tk2 | -0.043035144 | -0.02965296 | -0.054775567 | -0.05794236 | -0.071115849 | -0.029532131 | -0.033939322 | -0.023958392 | -0.009573138 | -4.16E-09 | -0.004273742 | -0.008578292 |
| d.n.tk3 | 0.039623903 | 0.058665659 | 0.032308495 | 0.029224822 | 0.023133531 | 0.044866898 | 0.036540341 | 0.055365867 | 0.04391471 | 0.06471655 | 0.071325091 | 0.067299165 |
| d.n.tpi | 0.088878291 | 0.056252531 | 0.074842577 | 0.073708284 | 0.11550973 | 0.049183964 | 0.102246586 | 0.030316698 | 0.016734766 | 0.016673682 | 0.039131826 | 0.045675599 |
| d.x.fum_b | 2.00E-16 | 1.84E-16 | -2.74E-16 | 2.56E-15 | 5.75E-01 | -1.11E-16 | 5.05E-01 | -9.36E-16 | 1.11E-17 | -5.22E-17 | -1.11E-15 | 3.77E-16 |
| f.n.BM | 0.075450635 | 0.080617211 | 0.079213585 | 0.07932364 | 0.085831193 | 0.066398943 | 0.062795489 | 0.070512068 | 0.044628507 | 0.053997909 | 0.061246851 | 0.06210774 |
| f.n.CO2_in | 3.00E-01 | 5.36E-01 | 5.040298261 | 3.790124991 | -1.48E-25 | 3.762109883 | 997.3113213 | 1.699027483 | 0.62056335 | 1.848483652 | -5.04E-16 | 2.91E-11 |
| f.n.Glc_in_1 | 0.822410745 | 0.82020645 | 0.825228293 | 0.82189351 | 0.824931144 | 0.832825994 | 0.830306201 | 0.831603315 | 0.823163684 | 0.819441493 | 0.822986715 | 0.822112599 |
| f.n.glk | 8.05E-02 | 9.01E-02 | 0.092373727 | 0.094124533 | 0.095775291 | 0.098047394 | 0.024907709 | 0.030147963 | 0.053801016 | 0.097856823 | 0.090343477 | 0.097609484 |
| f.n.gnd | 0.090695063 | 0.145891234 | 0.067177658 | 0.057906078 | 0.037227808 | 0.108895934 | 0.085246404 | 0.138626951 | 0.112557554 | 0.170935357 | 0.186441137 | 0.174406291 |
| f.n.kdgk | 3.61E-02 | 3.87E-02 | 0.036582528 | 0.036559476 | 0.038047692 | 0.036099672 | 0.037081471 | 0.043189382 | 0.036791297 | 0.037949615 | 0.038338513 | 0.038316157 |
| f.n.out_Kdg | 0.012139854 | 0.00469878 | 4.07E-21 | -1.83E-18 | 0.00026555 | 0.049726539 | 0.055476889 | 1.14E-18 | 0.080075943 | 0.060116819 | 0.059699459 | 0.054717327 |
| f.n.pdh | 1.037025761 | 1.103588638 | 1.136701894 | 1.061220193 | 1.079308469 | 1.140244396 | 1.047883933 | 1.229896039 | 1.190981391 | 1.215246809 | 1.104680396 | 1.125469726 |
| f.n.prot | 1.00E-05 | 1.00E-05 | 0.000349957 | 0.000307955 | 0.000256002 | 0.001867271 | 0.001898607 | 0.002338078 | 0.005161999 | 0.006244996 | 0.009545392 | 0.008799951 |
| f.n.pyc | 0.719935633 | 1.17E+00 | 0.526812873 | 0.555274372 | 0.894247221 | 0.490232099 | 1.079379193 | 0.94661892 | 0.689740715 | 0.67410465 | 0.951518062 | 1.049988079 |
| f.n.pyk | 0.471504941 | 0.630427478 | 0.555836334 | 0.536831006 | 0.521977786 | 0.586246694 | 0.50041088 | 0.665799009 | 0.745527286 | 0.773634003 | 0.81241144 | 0.8979951 |
| f.n.zwf | 0.147481824 | 0.155611961 | 0.125357747 | 0.119696853 | 0.142291227 | 0.146311664 | 0.114382979 | 0.074610781 | 0.093952296 | 0.166028362 | 0.181533424 | 0.186801981 |
| f.x.ald | 2.99E-16 | 3.80E-16 | 4.44E-16 | -6.66E-16 | 6.65E-15 | 2.22E-16 | 6.43E-02 | -3.99E-16 | 3.46E-18 | 4.25E-16 | -9.51E-17 | -3.89E-16 |
| f.x.eno | 0.231940706 | 2.71E-01 | 0.526515274 | 0.575752273 | 0.515582947 | 0.313177121 | 0.293539056 | 0.020399085 | 0.335413863 | 0.329664329 | 0.422720072 | 0.40001843 |
| f.x.fum_a | 2.50E-16 | 2.53E-16 | -2.29E-16 | 2.60E-15 | 5.75E-01 | -6.82E-17 | 5.05E-01 | -8.74E-16 | 6.24E-17 | -1.13E-18 | -1.05E-15 | 4.40E-16 |
| f.x.maldh | 0.103165461 | 0.596419537 | 0.343976905 | 0.18021322 | 0.021454401 | -4.59E-16 | 5.52E-16 | 1.79E-15 | -5.93E-16 | -2.17E-16 | -9.44E-16 | 3.33E-16 |
| f.x.pgi | 0.033409114 | 0.055362838 | 0.033597653 | 0.027076741 | 0.010988125 | 0.059677309 | 0.999 | 0.999 | 0.064887197 | 0.111690821 | 0.064082228 | 0.081784621 |
| f.x.pgk | -3.02E-17 | 6.73E-17 | 0.244621934 | 0.240659748 | 0.404977487 | 1.53E-01 | -2.12E-16 | 2.66E-15 | 0.395919109 | 0.230522397 | 0.285089399 | 0.273361599 |
| f.x.ta1 | 0.208421726 | 0.078538629 | 0.210570598 | 0.215672638 | 0.999 | 0.078092741 | 0.230077616 | -2.63E-16 | -9.37E-17 | 0.009413762 | 2.06E-16 | 0.012033916 |
| f.x.ta2 | 0.06277047 | 0.082848268 | 0.048304897 | 0.04985681 | 0.118146172 | 0.075405608 | 0.073761346 | 0.036653705 | 0.099414549 | 0.234821035 | 0.215477127 | 0.244203353 |
| f.x.tk1 | 0.226545943 | 0.434430351 | 0.276015607 | 0.189717684 | 0.124252366 | 0.269657701 | 0.348347038 | 0.249606783 | 0.788812896 | 0.270305984 | 0.48197712 | 0.328668738 |
| f.x.tk2 | 6.87E-17 | 6.43E-17 | 7.68E-17 | -2.54E-16 | 7.48E-16 | 2.44E-16 | 5.41E-17 | 5.58E-17 | 7.72E-17 | -2.41E-17 | -1.94E-16 | 2.78E-17 |
| f.x.tk3 | 0.003339059 | 0.016073889 | -3.71E-17 | 1.53E-16 | 0.03507381 | 0.004542671 | 3.56E-17 | 0.999 | 0.302686472 | 0.068546614 | 0.166567695 | 0.108161337 |
| f.x.tpi | 4.64E-16 | -9.95E-19 | -3.10E-17 | -3.79E-16 | 5.87E-01 | -6.66E-16 | 5.43E-01 | -1.39E-16 | 2.54E-18 | -1.92E-17 | -3.34E-16 | 2.28E-16 |
